# Amyloid-mediated RNA uptake by sperm for embryonic delivery

**DOI:** 10.64898/2025.12.10.693493

**Authors:** William S. Mallen, Nataliya Zabello, Susan Brenner-Morton, Simone Sidoli, Isabella D. Pirozzolo, Feng-Xia Liang, Joseph Sall, Jason Yin, Priyanka Mathews, Guoan Zhang, Carolyn L. Lee, Anthony W.P. Fitzpatrick

## Abstract

Seminal fluid RNA is essential for embryonic development, but the mechanism by which extracellular RNA is transferred from seminal fluid to sperm remains unclear. Here, we identify a role for functional amyloid fibrils in seminal fluid that bind and traffic RNA to mammalian sperm and, upon fertilization, the zygote. We show that highly cationic peptides in seminal fluid phase separate with RNA to form amyloid fibrils. Cryo-electron tomography reveals that these amyloid fibrils are internalized by sperm. Furthermore, biophysical and *in vitro* fertilization experiments show that amyloid-bound RNA reaches the zygote where molecular chaperones control release of RNA for translation. Our work uncovers a mechanism for RNA trafficking from seminal fluid to the zygote, establishing amyloid fibrils as potential tools for RNA delivery during fertilization.

---

Recent studies have shown that RNA trafficking from seminal fluid (SF) to developing gametes plays an epigenetic role in mammalian embryonic development (*1*, *2*). Prior research has demonstrated that environmental exposures, stress, and diet during adolescence can influence the next generation through sperm RNA (*3–5*). For example, male mice fed high-fat diets exhibit altered profiles of small RNA (sRNA) in their sperm compared to those on low-fat diets, leading to metabolic changes in their offspring (*4*). During sperm maturation in the epididymis, the sperm RNA payload undergoes dynamic changes between the caput and cauda regions through epididymal vesicles; however, the precise mechanism by which extracellular RNA is transferred from SF to spermatozoa remains unclear (*6*, *7*). In 2007, naturally occurring cationic amyloid fibrils were identified in SF and were found to enhance human-immunodeficiency virus (HIV) infectivity by neutralizing the virus’s negative charge and promoting viral entry into cells (*8*). In addition to their role in enhancing infectivity (*8*), SF amyloid fibrils may also serve beneficial functions such as either sperm selection or antimicrobial defense (*9*, *10*). Together, these findings raise the intriguing possibility that functional amyloid fibrils (*11*) in SF may bind and stabilize RNA, facilitate its uptake by sperm, and ultimately deliver it to the zygote upon fertilization (Fig. 1A). This proposed mechanism not only provides a route for RNA trafficking from SF to sperm but also highlights a potential therapeutic strategy for delivering exogenous nucleic acids to enable genetic modification for disease prevention.

**Fig. 1.**
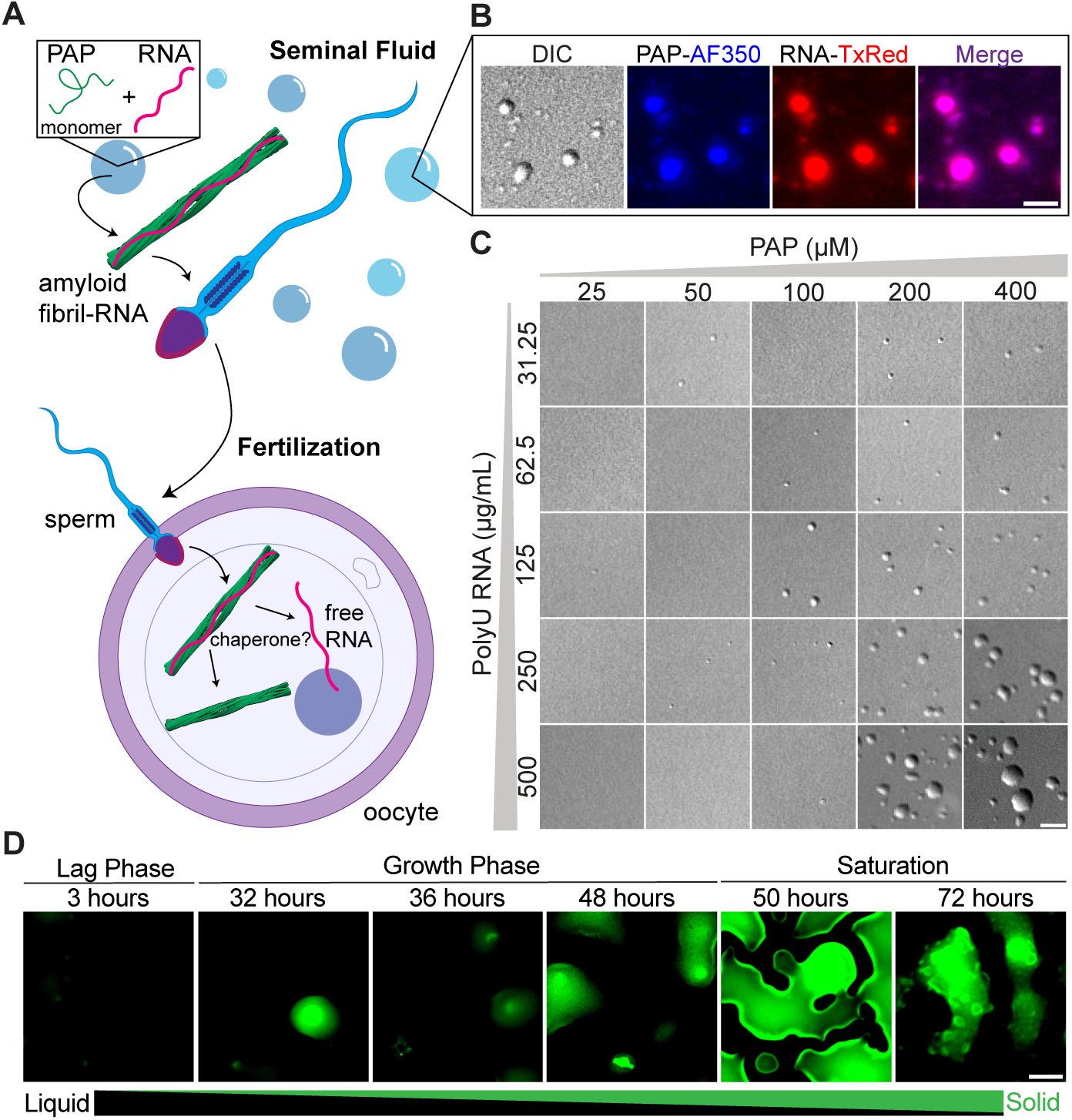
PAP phase separates with RNA to form amyloid fibrils. **(A)** Schematic of amyloid-mediated RNA transfer from seminal fluid to the embryo. PAP and RNA phase separate extracellularly, forming amyloid fibrils that are internalized by sperm and released into the zygote upon fertilization. RNA then dissociates from the fibril and can influence the genome. **(B)** Confocal microscopy of 400 µM PAP-AF350 with 500 µg/mL polyU RNA-TxRed showing phase separation and co-localization. Scale bar, 5 µm. **(C)** Phase separation of PAP with polyU RNA at varying concentrations. Representative images from 3 independent experiments (*n* = 3). Scale bar, 5 µm. **(D)** Time-course of PAP and 30-nt polyU RNA phase transition over 48 hours, monitored by ThT fluorescence. Scale bar, 5 µm.

## PAP phase separates with RNA

Peptides in semen known to form amyloid fibrils include fragments of prostatic acid phosphatase (PAP)—specifically residues 85-120 and 248-286—as well as semenogelin I and II (SEMG1 and SEMG2), which vary between residues 45-107 (*8*, *12*, *13*). While SEMG1 and SEMG2 fragments are evolutionarily conserved in primates (*12*, *14*), PAP fragments are conserved across all mammals, suggesting a potential role in reproduction (fig. S1). While SEMG1 and SEMG2 are secreted by seminal vesicles, PAP is secreted from the prostate into prostatic fluid and combines with fluid from seminal vesicles to form semen (*15*). Both PAP fragments are cationic; however, PAP(248-286) has the highest isoelectric point (pI = 10.21) and a net charge of +6.2 (*12*). Notably, PAP(248-286) was identified in the peptide fraction that exhibited the highest rate of HIV infectivity (*8*), demonstrating its capacity to bind negatively charged molecules such as RNA. Based on these characteristics, we selected PAP(248-286), which will now be referred to as PAP, to serve as a model to investigate the role of semen-derived amyloid fibrils.

As interactions between PAP and RNA have not previously been reported, we first examined the fragment’s primary sequence to identify regions that may promote RNA binding (fig. S2A). The abundance of positively charged residues and scarcity of negatively charged ones suggest a strong propensity for RNA interaction (*16–18*). Additionally, two intrinsically disordered regions (IDRs) are predicted within the fragment—between residues 248-257 and 283-286—with the former enriched in positively charged residues (fig. S2B) (*19*). The fragment also contains aggregation-prone regions that likely drive amyloid fibril formation (fig. S2C) (*20*). These disordered and aggregation-prone features are consistent with other semen-derived fragments (fig. S3, A-C). Given the presence of IDRs, we tested whether PAP undergoes liquid-liquid phase separation (LLPS). Although several residues are predicted to have LLPS potential, the overall low LLPS propensity score of -5 (*21*) suggests the fragment cannot phase separate on its own—a prediction we confirmed experimentally (fig. S3D).

To determine whether PAP coacervates with RNA, we fluorescently labeled the peptide with Alexa Fluor 350 and mixed it with polyU RNA tagged with Texas Red. Confocal microscopy revealed that PAP and RNA phase separate and co-localize in liquid droplets (Fig. 1B). This suggests that RNA promotes molecular crowding of PAP, resulting in local high concentrations of both peptide and RNA. In contrast, SEMG1(68–107) did not phase separate with polyU RNA, likely due to its lower net charge (fig. S3D). We observed that increasing concentrations of either PAP or RNA seemed to promote the formation of larger droplets (Fig. 1C), which was confirmed quantitatively (fig. S4A). To determine whether LLPS occurs under physiological salt conditions (as salt is known to disrupt LLPS (*16*)) we performed a NaCl gradient assay at constant peptide and RNA concentrations (fig. S4B). Given that prostatic and seminal fluid typically contain 100-200 mM NaCl (*22*, *23*), we found that PAP and RNA successfully phase separate within this range.

## RNA promotes amyloid fibril formation

Previous studies have demonstrated that synthetic PAP peptides can form amyloid fibrils independently when incubated at 37 °C under agitation (*8*, *24*). However, the mechanism by which these fibrils nucleate under physiological conditions remains unclear. Our biophysical experiments show that PAP undergoes LLPS in the presence of RNA, but whether, or not, LLPS leads to amyloid fibril formation is unknown. To investigate this potential liquid-to-solid phase transition, we performed an LLPS assay alongside a kinetic time-course experiment (Fig. 1D, fig. S5A). Thioflavin T (ThT), a dye that fluoresces upon binding to β-sheet structure at a wavelength of 482 nm, was added to a mixture of PAP and polyU RNA in two identical 96-well plates at room temperature with no agitation (*25*, *26*). Fluorescence intensity was measured over time in one plate, while confocal imaging was conducted on the other.

During the lag phase, phase separation was visible in differential interference contrast (DIC) microscopy, but no ThT fluorescence was observed, indicating the absence of amyloid fibrils (Fig. 1D). As the ThT signal increased between hours 32 and 48, corresponding to the growth phase (fig. S5A), fluorescence was observed from the droplets (Fig. 1D). By hour 72, the ThT signal on the kinetics curve plateaued, consistent with mature amyloid fibril formation (Fig. 1D; fig. S5A). Confocal imaging at this time revealed visible aggregates within what appeared to be solidified droplets (fig. S5A). These results suggest that LLPS precedes and precipitates the formation of PAP amyloid fibrils in the presence of RNA.

To confirm that our synthetic PAP amyloid fibrils made in the presence of RNA are indeed amyloid, cryo-electron microscopy (cryo-EM) revealed a predominantly flat, doublet (width ∼ 140 Å) fibril morphology with a helical rise of 4.7 Å—a molecular signature of amyloid structure (fig. S5, B and C). Moreover, we isolated the amyloid-containing sarkosyl-insoluble fraction from pooled human SF donors (table S1) and cryo-EM revealed a thinner, singlet (width ∼ 70 Å) amyloid fibril also with a helical rise of 4.7 Å (fig. S5, D and E).

## SF amyloid fibrils bind to various small RNAs

We next examined the types of RNA that could interact with PAP *in vitro* and *in vivo*. An LLPS assay shows RNA length influences droplet formation by mixing PAP with RNAs of varying nucleotide lengths. Longer sRNAs led to larger droplets (Fig. 2A, fig. S6A). Further LLPS assays using PAP with polyA, polyC, tRNA, and yeast RNA confirmed phase separation in all cases (fig. S6B). To further test whether, or not, RNA is essential for LLPS, we treated samples with RNase A, which cleaves the phosphodiester bond at the 3’ end of pyrimidine nucleotides (uracil and cytosine) (*27*). RNase A treatment disrupted LLPS in polyC and polyU RNA conditions (fig. S6B). However, droplets persisted in polyA RNA samples, while tRNA and yeast RNA conditions resulted in smaller droplets (fig. S6B). These findings indicate that RNA is critical for PAP-mediated LLPS, though droplet size may vary depending on RNA length and type.

**Fig. 2.**
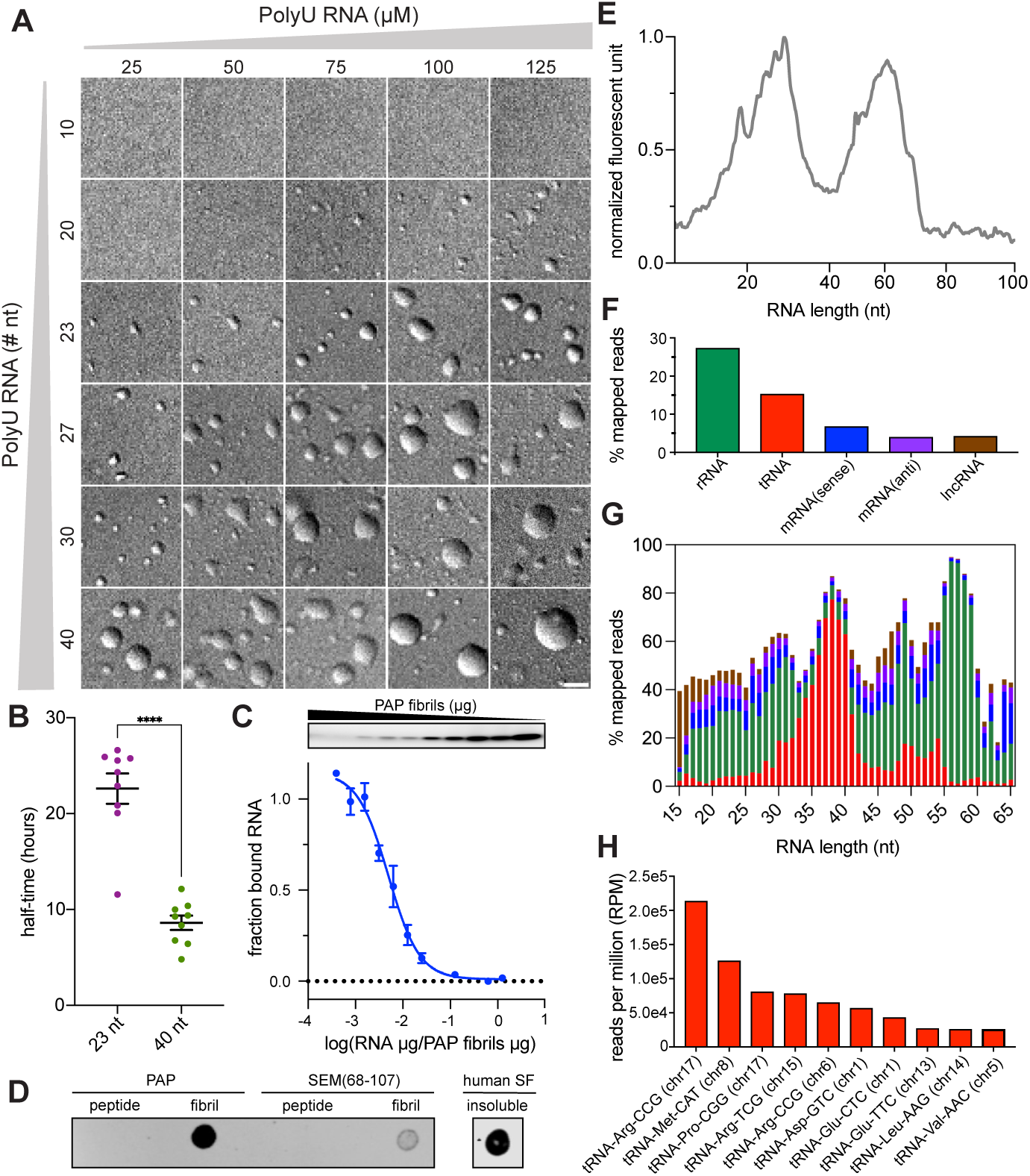
Semen-derived amyloid fibrils interact with diverse small RNAs. **(A)** Phase separation of PAP with polyU RNA at varying concentrations and nucleotide lengths. Representative images from 3 independent experiments (*n* = 3). Scale bar, 5 µm. **(B)** Half-time plot from kinetics curve in fig. S6E, showing the time at which 50% of PAP forms amyloid fibrils (n = 3). *P* < 0.0001 (****). **(C)** Gel electrophoresis showing unbound RNA (top) and corresponding binding curve (bottom) of RNA *per* PAP amyloid fibril. Solid line indicates sigmoidal fit; error bars represent standard error of the mean. Curve normalized to initial RNA amount (*n* = 3). **(D)** Dot plot detecting PAP amyloid fibrils using an in-house anti-PAP amyloid fibril IgM. **(E)** Size distribution and abundance of small RNA fragments in the amyloid-containing sarkosyl-insoluble fraction of cryopreserved human semen. **(F)** Percentage of mapped reads of the top 5 most abundant RNA species. **(G)** Percentage of mapped reads across RNA size distribution from RNA-sequencing analysis. See (F) for legend. **(H)** Reads per million (RPM) of the top ten tRNA derived fragments identified.

Kinetics show that polyU RNA significantly accelerates PAP amyloid fibril formation (fig. S6, C and D). As PAP amyloid fibrils do not form independently without agitation, kinetic assays were performed at 37 °C under agitation. The mean half-time (t_1/2_) for fibril formation was 26.0 ± 5.83 hours (fig. S6, C and D). With the addition of 40 µg/mL RNA, the mean half-time decreased to 7.96 ± 0.96 hours (fig. S6, C and D), indicating that RNA shortens the lag phase and promotes a faster transition from monomer to fibril. Moreover, the rate of fibril formation was dependent on RNA length. When PAP was incubated with either 23- or 40-nucleotide (nt) length polyU RNA, the mean half-times were 22.6 ± 4.74 hours and 8.61 ± 2.27 hours, respectively, suggesting longer sRNAs may present more nucleation sites (Fig. 2B, fig. S6E).

To explore how RNA interacts with PAP amyloid fibrils, we formed fibrils in the presence and absence of 30-nt polyU RNA-ATTO488. Fibrils formed without RNA were subsequently incubated with the labeled RNA. An electrophoretic mobility shift assay (EMSA) revealed a shift in RNA migration to higher molecular weights, indicating that RNA binds to PAP amyloid fibrils (fig. S7A). The smear observed in the gel reflects the heterogenous lengths of fibrils, each capable of interacting with RNA. These findings suggest that RNA can either be incorporated into fibrils during nucleation and growth or bind to preformed fibrils. We performed a sedimentation binding assay to determine the amount of synthetic PAP amyloid fibrils needed to bind all available RNA to reduce the likelihood of unbound RNA in future experiments (Fig. 2C). Fibrils were diluted at different concentrations and incubated with 30-nt polyU RNA-ATTO488. After centrifugation, supernatants were removed and analyzed by an EMSA to determine the amount of free (unbound) RNA. After quantification, a singlet PAP amyloid fibril captures at least 2 molecules of 30-nt polyU RNA-ATTO488.

To identify RNAs that endogenously interact with SF amyloid fibrils, we purified the sarkosyl-insoluble fraction from human semen (table S1). Mass spectrometry revealed PAP, SEMG1, and SEMG2 as among the most abundant proteins (fig. S7B; data S1). We also developed a conformation-specific IgM antibody against synthetic PAP amyloid fibrils (fig. S7C) and confirmed that intact PAP amyloid fibrils—not just peptides—are present in the purified sarkosyl-insoluble fraction by dot blot (Fig. 2D; table S2). To characterize the endogenous sRNAs associated with SF amyloid fibrils, we extracted RNAs <200 nucleotides in length from the sarkosyl-insoluble fraction of cryopreserved human semen from pooled donors (table S3). sRNA analysis revealed two prominent RNA size distributions: 10-40 nucleotides and 40-75 nucleotides (Fig. 2E). Due to PAP and RNA forming the largest droplets when using RNA between 30 and 40 nucleotides in our LLPS assay, we were most interested in which RNA species are enriched at that size. RNA-sequencing and analysis with sRNAbench (*28*) showed that the majority of sRNAs were rRNA and tRNA derived fragments (tDFs) (Fig. 2F). The first peak was predominantly tDFs, while the second consisted mostly of rRNA fragments (Fig. 2G). Specifically, tRNA-Arg-CCG, tRNA-Met-CAT, and tRNA-Pro-CGG were the top three tRNAs identified (Fig. 2H). Notably, tDFs have been shown to alter gene expression and pass on epigenetic traits to offspring (*29*). Other RNAs detected to a lesser extent included long non-coding RNA, mRNA, microRNA (miRNA), yRNA, and small nuclear RNA. Interestingly, the supernatant of a sarkosyl-insoluble pellet contained only 0.01% of tRNA, suggesting that the majority of tDFs are enriched in the amyloid-containing sarkosyl-insoluble fraction (fig. S8). Having confirmed the presence of amyloid fibrils in semen and their association with RNA, we next examined whether these fibrils can be internalized by mammalian spermatozoa.

## Cryo-ET identifies amyloid fibril uptake in spermatozoa

Cryo-electron tomography (cryo-ET) was used to determine whether mammalian spermatozoa internalize PAP amyloid fibrils. Bovine sperm was selected as a model due to its large cell size. Synthetic PAP amyloid fibrils were covalently linked to 3 nm DAPI-labeled gold nanobeads by a stable Au-N bond formed between the gold surface and lysine side chains (*30*). Mature spermatozoa were incubated with DAPI-gold-PAP amyloid fibrils and stained with propidium iodide (PI) and SYBR14 to distinguish live cells with intact membranes (Fig. 3A) (*31–33*). Sperm were sorted by fluorescence-activated cell sorting (FACS) on live cells (Fig. 3B; fig. S9), which were then frozen on 200-mesh hexagonal grids (Fig. 3C, top) (*34*). Cryo-correlative light and electron microscopy (cryo-CLEM) was used to identify sperm cells potentially containing internalized fibrils based on DAPI fluorescence (Fig. 3C, bottom). Selected cells were then milled using cryo-focused ion beam scanning electron microscopy (cryoFIB-SEM) to create lamellae approximately 20 µm long and 15 µm wide (Fig. 3D). Overlaying a cryo-CLEM image with a lamella indicated the presence of DAPI-labeled fibrils in the midpiece of a sperm cell (Fig. 3E).

**Fig. 3.**
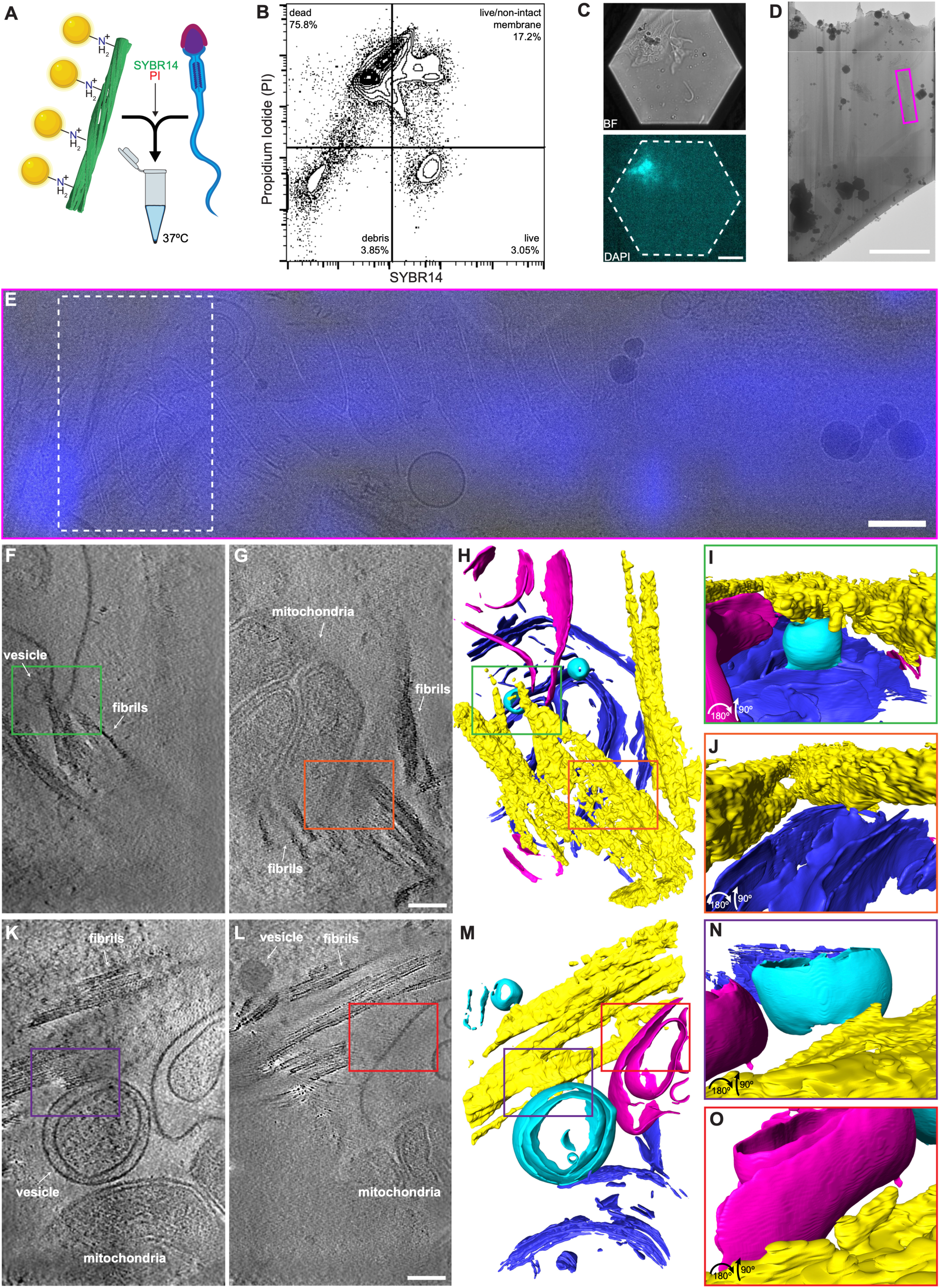
Cryo-ET reveals internalization of PAP amyloid fibrils by sperm. **(A)** Schematic of dairy sperm preparation for cryo-electron tomography (cryo-ET). Amyloid fibrils covalently linked to 3 nm gold nanobeads were incubated with sperm (50 µg/mL) and later stained with SYBR14 and PI. **(B)** FACS gating on SYBR14+ PI-sperm to isolate single cells for analysis. **(C)** Cryo-CLEM image of sperm frozen on 200-mesh hexagonal Au grids. DAPI fluorescence marks PAP amyloid fibrils linked to gold nanobeads. Scale bar, 20 µm. **(D)** Montaged cryo-TEM image of sample lamella. Scale bar, 1.5 µm. **(E)** Enlarged region of pink rectangle in (D) overlaid with DAPI channel from cryo-CLEM. Scale bar, 250 nm. **(F)** Averaged top slice of a tomographic volume from the white dashed box in (E). **(G)** Averaged bottom slice of the same tomographic volume in (E). Scale bar, 100 nm. **(H)** Segmentation of the tomogram reconstructed from the tilt series in (E). Color code: blue, mitochondria; yellow, amyloid fibrils; cyan, vesicles; pink, unidentified membrane. **(I)** Zoomed-in view of the area outlined in the green box in (F) and (H). **(J)** Zoomed-in view of the area outlined in the orange box in (G) and (H). **(K)** Averaged top slice of a second tomographic volume. **(L)** Averaged bottom slice of the same volume. Scale bar, 100 nm. **(M)** Segmentation of the tomogram from (K) and (L). Same color code as in (H). **(N)** Zoomed-in view of the area outlined in the purple box in (K) and (M). **(O)** Zoomed-in view of the area outlined in the red box in (L) and (M).

Reconstructed tomograms revealed the presence of amyloid fibrils, mitochondria, vesicles, and unidentified membranes in the sperm midpiece (Fig. 3, F and G). PAP amyloid fibrils were identified by their decoration with gold nanobeads. In one tomogram (Fig. 3, F and G), segmentation revealed direct interactions between fibrils and cellular components (Fig. 3H; movie S1). An averaged *z*-slice shows a fibril in close proximity to a vesicle and an unidentified membrane (Fig. 3, F and I), and segmentation confirmed direct contact. Another averaged *z*-slice shows fibrils pressing against the outer mitochondrial membrane, also confirmed by segmentation (Fig. 3, G and J). A second tomogram revealed PAP amyloid fibrils located within a sperm’s residual cytoplasmic droplet (Fig. 3, K-M; movie S2). Again, segmentation confirmed contacts between fibrils and vesicles (Fig. 3, K and N) as well as unidentified membranes (Fig. 3, L and O). While previous studies visualized the plasma membrane of mouse sperm using thicker (∼300 nm) lamellae (*35*, *36*), our thinner 150 nm sections did not clearly resolve the plasma membrane in bovine sperm. To confirm fibril internalization, we employed high pressure freezing-freeze substitution serial sectioning electron tomography (HPF-FS ssET).

To visualize the amyloid fibrils at low magnification, the 3 nm gold nanobeads underwent silver enhancement to enlarge to ∼ 20 nm. The beads were visible by HPF-FS ssET and were confirmed to be internalized in some *z*-slices. In 3D tomograms, the plasma membrane was resolved (Fig. 4). An averaged *z*-slice revealed multiple internalized fibrils within a sperm cell (Fig. 4A). In the midpiece, two fibril clusters were observed in close contact with axonemes (Fig. 4, B and C), and tomograms showed fibrils appearing and disappearing within the midpiece (movie S3). Segmentation revealed interactions between PAP amyloid fibrils and the plasma membrane, possibly being internalized (Fig. 4C; movie S4). Farther along the midpiece, fibrils were also found in contact with axonemes near a cytoplasmic droplet (Fig. 4, D and E; movie S5). Finally, fibrils were observed near the sperm head, in proximity to the nucleus (Fig. 4, F and G; movie S6).

**Fig. 4.**
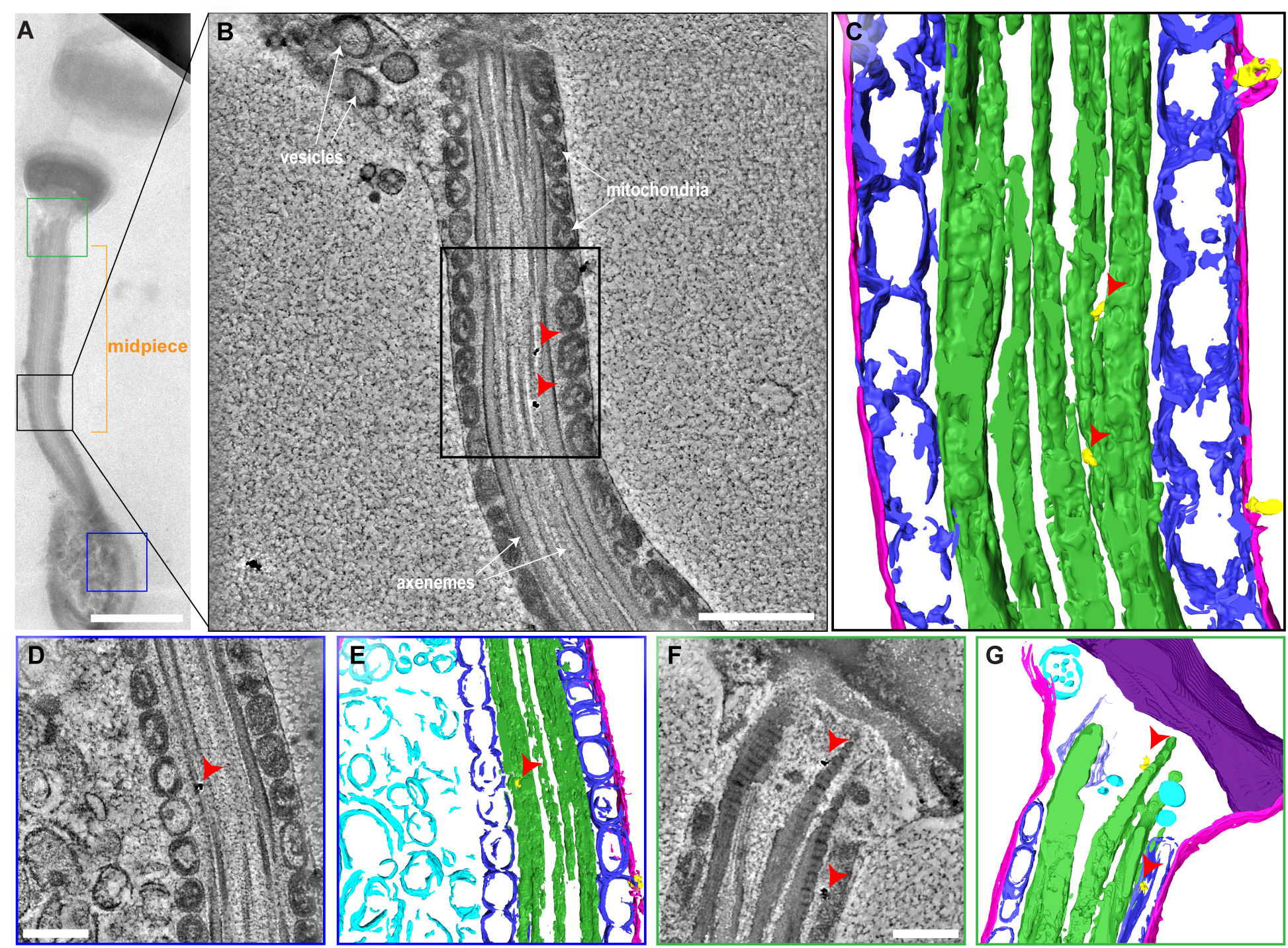
Volume EM identifies regions of PAP amyloid fibril internalization in sperm. **(A)** Averages of TEM images of bovine sperm. Scale bar, 2 µm. **(B)** Central slice of a tomographic volume from a dual-axis tilt series targeting the black-boxed region in (A). Red arrows indicate gold nanobeads. Scale bar, 500 nm. **(C)** Segmented zoomed-in view of the black-boxed region in (B). Color code: pink, plasma membrane; blue, mitochondria; green, axoneme; yellow, amyloid fibrils. Red arrows indicate gold nanobeads. **(D)** Central slice of a tomographic volume from a dual-axis tilt series targeting the blue-boxed region in (A). Red arrows indicate gold nanobeads. Scale bar, 250 nm. **(E)** Segmented tomogram of (D). Color code same as (C); cyan, unidentified vesicles. Red arrows indicate gold nanobeads. **(F)** Central slice of a tomographic volume from a dual-axis tilt series targeting the green-boxed region in (A). Red arrows indicate gold nanobeads. Scale bar, 250 nm. **(G)** Segmented tomogram of (F). Color code same as (E); purple, nucleus; light purple, centriole. Red arrows indicate gold nanobeads.

Although cryo-ET and HPF-FS ssET confirmed that PAP amyloid fibrils are internalized by bovine sperm, the mechanism of entry and the proportion of cells that internalize fibrils remain unclear. A previous study has shown that PAP amyloid fibril uptake by TZM-bI cells is not mediated by phagocytosis (*8*). To investigate whether receptor-mediated endocytosis is involved (*37*), human sperm isolated from cryopreserved semen was treated with 0.25% trypsin for 3 minutes (table S4) (*38*). The cells were then incubated with Alexa Fluor 350-labeled PAP amyloid fibrils and sorted on live cells. FACS analysis revealed that 2.44 ± 0.18% of live sperm internalized fibrils (Fig. 5A, top). After trypsin treatment, this percentage dropped to 0.30 ± 0.20% (Fig. 5A, bottom; fig. S10), suggesting that internalization is largely receptor-mediated and dependent on surface proteins cleaved by trypsin. However, the efficiency (Fig. 5B) is an underestimate as the mature sperm used were cryopreserved, rather than fresh, and the fibrils were heavily fluorescently stained, possibly occluding receptor-binding sites.

**Fig. 5.**
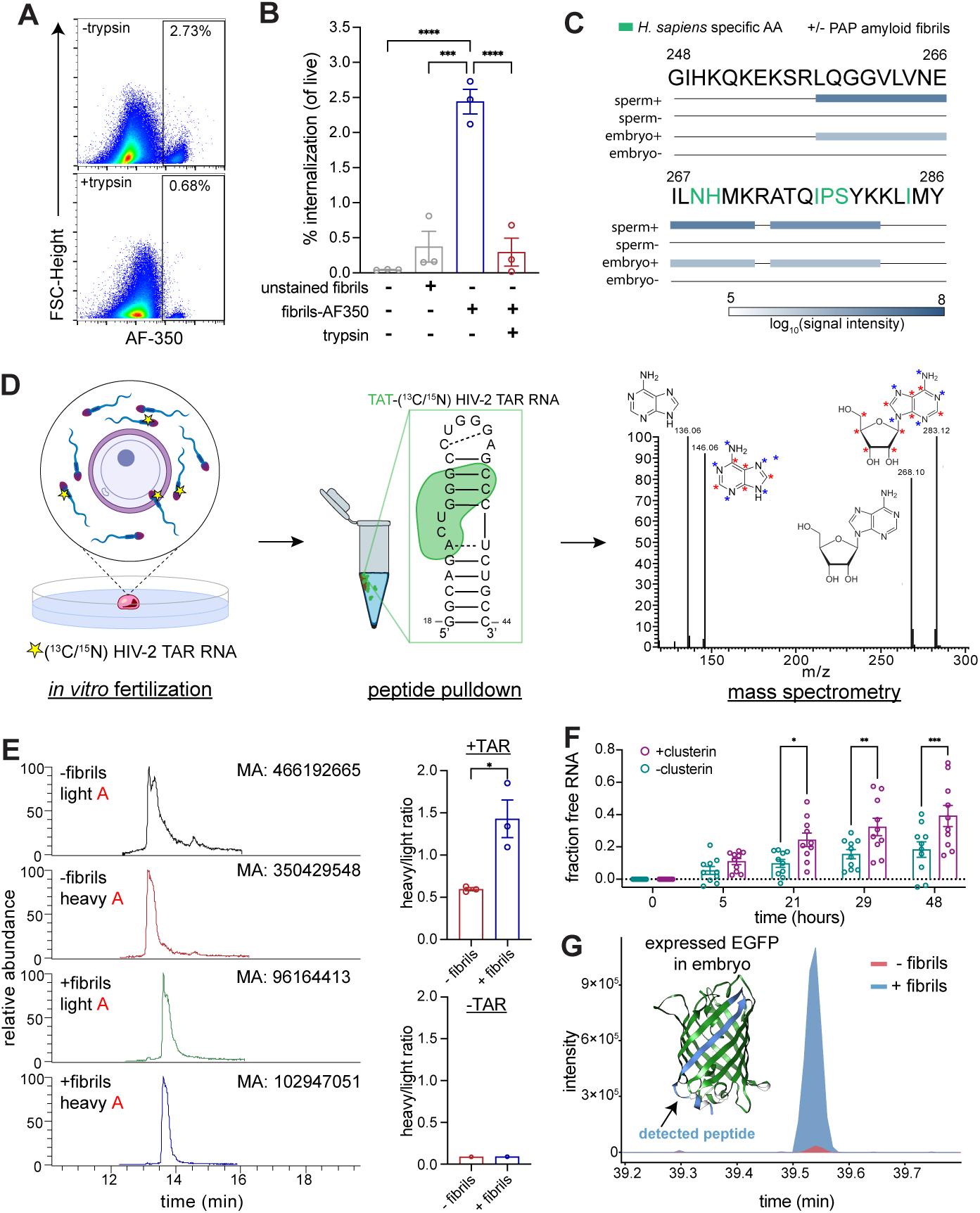
PAP amyloid fibrils and RNA are tracked from sperm to zygote. **(A)** FACS showing percentage of sperm cells gated for AF-350 in 1 representative trial. **(B)** Percent internalization of AF-350 labeled PAP amyloid fibrils by human sperm (*n* = 3). *P* values: 0.0001 (***); <0.0001 (****). **(C)** Mass spectrometry detection and intensity of *H. sapiens* PAP peptides in murine sperm and embryos from IVF. Two peptide fragments were detected: residues 258-272 and residues 274-281. Green lettering denotes human specific amino acids in PAP(248-286). **(D)** Schematic of murine IVF using isotopically labeled (^13^C/^15^N) HIV-2 TAR RNA. Red asterisks indicate ^13^C and blue asterisks indicate ^15^N labeling on adenine and adenosine. Strategy for enrichment and mass spectrometry detection is shown. **(E)** Representative mass spectrometry chromatograms showing retention times of light and heavy adenosine prior to fragmentation (left) and heavy adenine abundance relative to unlabeled adenine in the presence (*n* = 3) or absence (*n* = 1) of TAR RNA and PAP amyloid fibrils (right). MA: mass abundance. *P* = 0.0204 (*). **(F)** RNA dissociation assay showing fraction of RNA released from PAP amyloid fibrils over 48 hours, in the presence or absence of 50 µg clusterin (*n* = 10). *P* values: 0.0316 (*); 0.0089 (**); 0.0007 (***). **(G)** Total intensity of EGFP peptide detected in mass spectrometry. EGFP molecule shows region (light blue) that was detected.

## PAP amyloid fibrils mediate sperm uptake and zygotic delivery of RNA

While cryo-ET and FACS show internalization of PAP amyloid fibrils (Figs. 3 and 4), whether bound RNA is ultimately delivered to the zygote upon fertilization remained an outstanding question. To investigate, we first tracked PAP amyloid fibrils to see if they are present in the embryo by performing murine *in vitro* fertilization (IVF). Fresh C57BL/6 mouse sperm were incubated in the presence or absence of PAP amyloid fibrils prior to IVF. Following fertilization, mass spectrometry was performed on the protein lysates of the murine sperm and the resulting 2-cell stage embryos from IVF. Analysis detected two human, not mouse, PAP fragments spanning residues 258-272 and 274-281 in murine sperm and embryos (Fig. 5C). Although the precise mechanism by which amyloid fibrils are transferred from sperm to embryo remains unknown, it is plausible that they follow a pathway similar to that of paternal mitochondria—transferred at fertilization and subsequently targeted for degradation during embryonic development (*39*).

To track amyloid-mediated RNA transfer from sperm to the zygote, we performed IVF using readily purified, isotopically labeled (^13^C/^15^N) HIV-2 trans-activation response (TAR) RNA. Total RNA was extracted from 2-cell embryos, and TAR RNA was enriched by affinity capture using a biotinylated Tat(47–57) peptide (TAR-binding domain) immobilized on streptavidin-coated magnetic beads (Fig. 5D) (*40*, *41*). The enriched fractions were analyzed by mass spectrometry to quantify heavy adenine abundance relative to that of unlabeled adenine, the fragmented portion of adenosine (Fig. 5E). The presence of PAP amyloid fibrils significantly increased TAR RNA transfer to the embryo (Fig. 5E), indicating that PAP amyloid fibrils enhance RNA delivery from seminal fluid to sperm and ultimately to the zygote. This led us to ask: how does RNA dissociate from PAP amyloid fibrils?

Mass spectrometry analysis of the insoluble fraction from human SF revealed an abundance of molecular chaperones, such as clusterin and heat shock cognate 70 (Hsc70) (fig. S7B; data 1). Clusterin has been previously shown to disaggregate PAP amyloid fibrils (*42*). To test whether clusterin can dissociate RNA from PAP amyloid fibrils, we performed a dissociation assay in which preformed fibrils made in the presence of polyU RNA-ATTO488 were incubated with or without clusterin at 37 °C with no agitation. Aliquots were taken over 48 hours, centrifuged, and supernatants removed for analysis. An EMSA was performed on the supernatants to determine the quantity of free RNA, which indicates dissociated RNA (fig. S11A). Clusterin significantly dissociated RNA from PAP over the course of 48 hours compared to fibrils with no clusterin (Fig. 5F; fig. S11B). The rate of fibril dissociation was determined to be 1.05 × 10^-2^ ± 8.17 × 10^-4^ fraction bound RNA *per* hour in the presence of clusterin and 5.40 × 10^-3^ ± 4.50 × 10^-4^ fraction bound RNA *per* hour without clusterin. Additionally, mass spectrometry of murine sperm and embryos confirmed the presence of mouse clusterin (data S2). These findings suggest that the amyloid-RNA complex could be interacting with clusterin in SF, sperm, and the embryo.

To demonstrate that fibril trafficked RNA is translationally active in the embryo, IVF was performed again, but instead with enhanced green fluorescent protein (EGFP) mRNA. Total embryonic RNA was extracted and amplified by template switching RT-PCR. Amplified nested EGFP cDNA bands were observed on an agarose gel (fig. S12). As the presence of EGFP mRNA was detected, mass spectrometry was performed on the protein lysates of 2-cell stage embryos that were fertilized by sperm incubated with EGFP mRNA in the presence or absence of PAP amyloid fibrils. While EGFP protein was detected in both samples, the expression level was increased in the embryos that were fertilized by sperm incubated in the presence of PAP amyloid fibrils by at least an order of magnitude (Fig. 5G). This data confirms that RNA trafficked to the embryo by PAP amyloid fibrils is translationally active.

## Mechanism of amyloid-mediated RNA transfer

While the role of paternal RNA in embryonic development and epigenetic inheritance is well established (*3*, *7*, *43–45*), our work identifies semen-derived amyloid fibrils as previously unrecognized extracellular carriers of RNA that can bridge SF, sperm, and the early embryo. By combining biophysical, proteomic, imaging, RNA sequencing, and functional delivery assays, we show that (i) a semen-derived, cationic, amyloidogenic peptide, PAP, coacervates with RNA into phase separated condensates that mature into amyloid fibrils (Fig. 1), (ii) endogenous semen-derived amyloid fractions are enriched for tDFs (Fig. 2), and (iii) amyloid-RNA complexes can be internalized by sperm (Figs. 3 and 4) and transferred to embryos for translation of exogenous proteins at fertilization (Fig. 5). Given that it takes sperm anywhere from 15-45 minutes to hours and days to fertilize an oocyte in the female reproductive tract, amyloid-mediated RNA transfer is physiologically relevant (*46*). These findings reveal a conceptual link between functional amyloids in semen and sperm RNA biology, while at the same time establishing an amyloid-based platform for RNA delivery at fertilization.

A central mechanistic insight is that RNA can act upstream of SF amyloid fibril formation by concentrating amyloidogenic peptides into a phase-separated state. *In vitro*, PAP coacervates with RNA into liquid droplets under physiologically relevant salt conditions (Fig. 1C, fig. S4). These droplets transition over time into ThT-positive assemblies, consistent with a liquid-to-solid maturation process (Fig. 1D, fig. S5A). RNA also accelerates fibrillization kinetics (fig. S6, C and D), and RNA length reduces the corresponding half-times (Fig. 2B). Together, these observations suggest a framework in which electrostatically driven condensation provides a nucleation mechanism for PAP amyloid fibril formation, transitioning from a dynamic extracellular peptide-RNA mixture into a stable, transportable amyloid-RNA complex.

Sarkosyl-insoluble fractions from human SF contain a bimodal distribution of sRNAs (Fig. 2E) that is dominated by tRNA- and rRNA-derived fragments, along with miRNAs and other noncoding RNAs (Fig. 2, F-H). The co-enrichment of PAP amyloid fibrils and molecular chaperones, such as clusterin and Hsc70 (fig. S7B), suggests that the SF amyloid-RNA complex is a dynamic, regulated structure *in vivo*. Our observation that clusterin can accelerate the disassembly of PAP amyloid fibrils and release bound RNA (Fig. 5F) demonstrates that the amyloid-RNA state is not irreversible, but instead one that can be modulated by chaperone-regulated disassembly, potentially enabling controlled exposure of sperm and the embryo to RNA over time.

Internalization of the amyloid-RNA complex into sperm is supported by direct structural and cell-based evidence that SF amyloid fibrils can traverse the plasma membrane. Cryo-ET visualizes gold-decorated PAP amyloid fibrils inside bovine sperm with segmented contacts to vesicles, unidentified membranes, and outer mitochondrial membranes (Fig. 3, F-O). Complementary HPF-FS ssET resolves fibril networks within the midpiece, including near axonemes and near the sperm head in proximity to the nucleus (Fig. 4). In human sperm, uptake is heterogeneous and strongly reduced by brief trypsinization, consistent with receptor-mediated internalization (Fig. 5A). This heterogeneity raises the possibility that amyloid-mediated uptake is regulated, potentially coupling the sperm surface state to the acquisition of extracellular RNA cargo *via* this route.

Our IVF experiments are the first demonstrated use of amyloid fibrils for nucleic acid delivery to the embryo for translation of exogenous proteins (Fig. 5, E and G). Endogenously, the sRNAs, such as tDFs, that interact with SF amyloid fibrils most likely contribute to gene silencing and regulatory mechanisms (Fig. 2, F-H). Unlike virus-like particles (VLPs)—such as Gag and Gag-like proteins—which typically bind nucleic acids containing specific sequence motifs, PAP amyloid fibrils can associate with a wide range of RNA molecules through nonspecific electrostatic interactions (Fig. 2, E-H) (*47–49*). VLPs, unless specifically engineered, generally package their own mRNA and are assembled intracellularly prior to exocytosis (*50*). Moreover, most endogenous Gag-like proteins in humans have lost the nucleocapsid domain, the principal RNA-binding region found in canonical Gag proteins (*49*). In contrast, PAP amyloid fibrils present a positively charged, extended surface area that promotes nonspecific, extracellular RNA binding and stabilization. Our tomograms also suggest that PAP amyloid fibrils cluster (Fig. 3, K and L), potentially increasing RNA binding and shielding RNA from the surrounding environment. This characteristic is particularly advantageous in harsh conditions, such as the acidic environment of the female reproductive tract (*9*). Additionally, cells may regulate their response to amyloid-trafficked RNA by controlling the timing of amyloid-RNA fibril disassembly by chaperones.

Current gene therapy approaches using adeno-associated viruses (AAVs) carry risks, particularly because the viral vector can trigger immune recognition and inflammation. For example, two AAV-based therapies are now approved for severe hemophilia A and B, but even in these cases, immune responses and dose-related toxicities remain concerns (*51*). In contrast, SF amyloid fibrils could theoretically enable delivery of therapeutic genetic material at fertilization, offering a non-viral strategy for families with a high predicted risk of transmitting an X-linked disorder.

Realizing the full biological and technological scope of SF amyloid fibrils will require defining the entry mechanism and its determinants on sperm surfaces, establishing the intracellular localization and fate of delivered RNAs with live-cell microscopy, and testing whether endogenous SF amyloid-associated RNA measurably influences early embryonic gene regulation by translational control or gene silencing. Nevertheless, our results identify a novel amyloid-based mechanism for extracellular RNA trafficking to the germline and provide a foundation for designing fibrillar delivery vehicles for RNA transport or therapeutic genetic modification—including potential applications in embryonic gene editing for disease prevention.

## Acknowledgements

The authors are grateful to Drs. S. Lomvardas, O.J. Rando, S. Tavazoie, R.S. Mann, and A.N. Chang for helpful discussions. We would also like to thank Humberto Ibarra from Cellular Imaging at Columbia University Zuckerman Institute, R. Grassucci and Dr. Z. Zhang for help collecting single-particle data at the Columbia University Cryo-Electron Microscopy Center including on the Titan Krios housed at Zuckerman Institute, the Ho lab for advice and protocols on cryoFIB-SEM and cryo-ET, and Ira Schieren and Max Wallach from Flow Cytometry at Columbia University Zuckerman Institute.

## Author contributions

A.W.P.F. and W.S.M. conceived the experiments. W.S.M., A.W.P.F., and I.D.P. performed RNA-seq and analysis. W.S.M., A.W.P.F., and N.Z. performed IVF. W.S.M., A.W.F.P., and S.S. performed and analyzed RNA mass spectrometry experiments. W.S.M., A.W.F.P., and G.Z. performed and analyzed protein mass spectrometry experiments. W.S.M., A.W.P.F., F.L., J.S., and J.Y. performed HPF-FS ssET. W.S.M., A.W.P.F., C.L., and S.B. developed antibody. W.S.M., A.W.P.F., and P.M. prepared phylogenetic trees. W.S.M. and A.W.P.F. performed cryo-EM, cryo-ET, and all other experiments. W.S.M. and A.W.P.F. prepared the manuscript, and A.W.P.F. supervised the project.

## Data and materials availability

All data are available in the manuscript or the supplementary materials.

## Materials and Methods

### Liquid-liquid phase separation assays

For all LLPS assays, PAP(248-286) peptides (Anaspec) were mixed with RNA variants in LLPS buffer (25 mM HEPES, 150 mM NaCl, pH 7.4) and immediately pipetted onto microscopic slides (FisherScientific) with cover glass (FisherScientific). DIC imaging was done with an inverted A1R confocal microscope (Nikon) at 60× magnification. For fluorescent LLPS assays, PAP(248-286) peptides were labeled with Alexa Fluor 350 (Thermo Fisher Scientific) prior to mixing with 30-nucleotide polyU RNA tagged with Texas Red (IDT) and imaged for fluorescence (360-20/455-20 and 620-30/700-35 filters). For salt concentration gradient LLPS assays, NaCl concentrations in LLPS buffer were adjusted accordingly. Lastly, for RNase A assays, 500 µg/mL RNase A was added to each condition after PAP(248-286) and RNA phase separation.

### Fibril formation assays

PAP(248-286) peptides (Anaspec) at 2 mg/mL, RNA variants such as polyU RNA at 40 µg/mL (non-length specific) or 30 µM (length specific), 40 µM ThT (MedChemExpress), and kinetics buffer (50 mM sodium phosphate monobasic monohydrate, 150 mM NaCl, pH 7.4) were mixed and plated into a Costar 96-well plate (Corning) with clear bottom. Plates were incubated at 37 °C at 700 rpm in double-orbital mode and monitored for ThT fluorescence (440-10/480-10 filters) every 5 minutes in a FLUOstar Omega microplate reader (BMG Labtech).

### EMSAs

PAP(248-286) peptides (Anaspec) at 2 mg/mL were incubated with and without 30-nucleotide polyU RNA-ATTO488 (IDT) in kinetics buffer in an Eppendorf thermomixer at 37 °C and 1400 rpm for 24 hours. Once synthetic PAP(248-286) amyloid fibrils have formed, samples were mixed with a non-SDS gel purple loading dye (NEB) and loaded onto a Novex 4-20%, Tris-Glycine Plus WedgeWell Gel (Invitrogen). Samples were run with 0.5× TBE buffer (Thermo Fisher Scientific) at 120-volts. Gel was then rinsed with water and imaged on a Typhoon FLA 9500.

### Sedimentation binding assay

PAP(248-286) amyloid fibrils were prepared with 2 mg/mL of monomer. Fibrils were centrifuged at 21,130 × *g*, 30 min, 4 °C and supernatant was removed. The amount of protein in the supernatant detected by Nanodrop 8000 (Thermo Scientific) was subtracted from the total PAP(248-286) monomer used to determine the amount of fibrils in the pellet. Kinetics buffer was added to the pellet to bring the concentration to 2 mg/mL. Fibrils were diluted in kinetics buffer by the following: 1:3.125, 1:6.25, 1:12.5, 1:25, 1:50, 1:100, 1:200, 1:1000, 1:5000, and 1:10,000. Each concentration of PAP(248-286) amyloid fibrils was incubated with 100 nM 30-nucleotide polyU RNA-ATTO488 in 50 mM HEPES, 150 mM NaCl, pH 7.4. Samples were centrifuged at 21,130 × *g* for 30 minutes at 4 °C. Supernatant was removed and loaded onto a Novex™ tris-glycine, 4-20% WedgeWell (Thermo Fisher Scientific) at 120 V with non-SDS loading dye. The gels were imaged on a Typhoon FLA 9500 instrument and gel intensities were quantified in Fiji.

640 ng PAP amyloid fibrils pulls down 1 ng RNA:

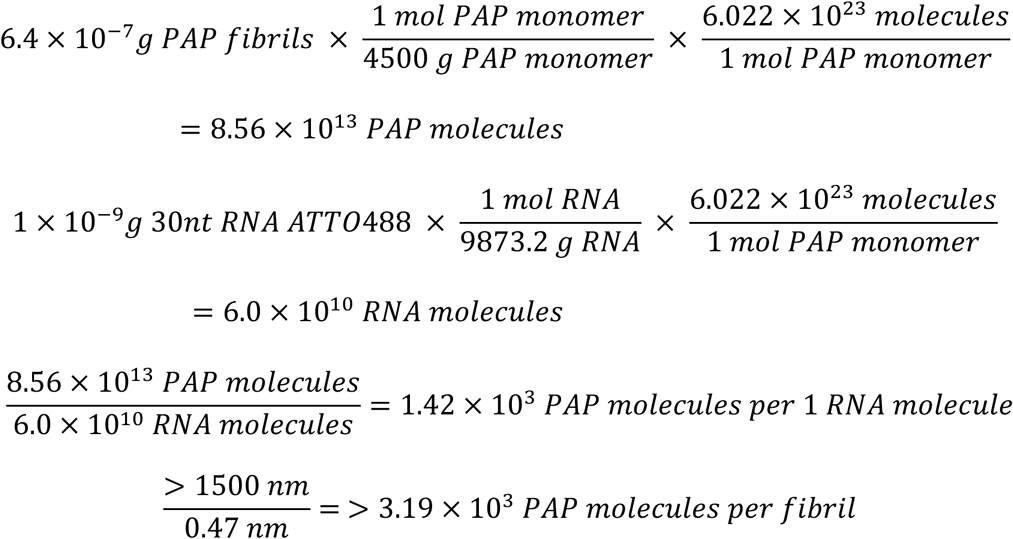

### Phase transition assays

PAP(248-286) peptides (Anaspec) at 2 mg/mL were incubated with 60 µg/mL polyU RNA (Sigma) and 40 µM ThT (MedChemExpress) in two identical 96-well glass bottom plates (Cellvis) in kinetics buffer at RT. Imaging of one plate was performed at various time points over the course of 72 hours using an inverted A1R confocal microscope (Nikon) at 60× magnification monitoring for fluorescence (480-15/535-20 filters). Meanwhile, ThT fluorescence was observed in the Omega microplate reader as described previously.

### RNA disassociation assay

PAP(248-286) amyloid fibrils were made in the presence of 2 mg/mL PAP peptides (Anaspec) and 30 µg/mL 30-nucleotide polyU RNA-ATTO488 in 200 mM phosphate buffer, pH 7.2, at 37 °C and 700 rpm for 48 hours. After formation, fibrils were washed and resuspended in 500 µL buffer and 50 µg clusterin (Acro Biosystems). The sample was incubated at 37 °C with no shaking and 30 µL aliquots were taken at 0, 5, 21, 29, and 48 hours and immediately snap-freezed in liquid nitrogen. After 48 hours, samples were thawed and centrifuged at 21,130 × *g*, 30 min, 4 °C. Supernatant was removed into a separate tube. Supernatants were then loaded on a Novex™ tris-glycine, 4-20% WedgeWell (Thermo Fisher Scientific) with non-SDS loading dye at 120 V. Gels were imaged on a Typhoon FLA 9500 instrument and intensities of the bands were measured in Fiji. “Fraction free RNA” is band intensity as a fraction of control band intensity.

### Isolation of sarkosyl-insoluble fraction from human SF

Seminal fluid (Lee Biosolutions) was thawed in 37 °C water bath and centrifuged at 400 × *g* and 4 °C for 10 minutes. The supernatant was transferred to centrifuge tubes (Beckman Coulter) and topped off with Tris buffer (10mM Tris pH 7.4, 0.8 M NaCl, 1 mM EGTA, 5 mM EDTA, and 10% sucrose). Tubes were centrifuged at 100,000 × *g* for 1 hour at 4 °C. The pellet was resuspended in Tris buffer and 2% sarkosyl. Sample rotated overnight at 4 °C and then centrifuged at 4,000 × *g* for 10 minutes at 4 °C. Supernatant was isolated and centrifuged again at 100,000 × *g* as before. Pellet was resuspended in freezing buffer (20 mM Tris and 150 mM NaCl) for cryo-EM.

### RNA extraction, library preparation, and sequencing

Cryopreserved human semen (Lee Biosolutions) was centrifuged at 500 × *g* for 5 minutes at 4 °C and supernatant was transferred into a centrifuge tube (Beckman Coulter) with SUPERase•In^TM^ RNase Inhibitor (Thermo Fisher Scientific). The sample was centrifuged at 400,000 × *g* at 4 °C for 1 hour. The supernatant was removed, and the pellet was resuspended in Tris buffer, 2% sarkosyl, and more RNase inhibitor. The sample was incubated at 4 °C while rotating gently. After 1 hour, the sample was centrifuged at 500 ×*g* for 10 minutes at 4 °C. The supernatant was transferred to a centrifuge tube and centrifuged at 400,000 × *g* again for 1 hour. The supernatant was removed, and pellet was immediately resuspended in RLT buffer for RNA extraction using the RNeasy Micro Kit (Qiagen) isolating fractions <200 nucleotides. RNA quality was assessed on a 2100 Bioanalyzer (Agilent). A smRNA cDNA library was prepared from isolated RNA by using SMARTer® smRNA-Seq Kit for Illumina® (Takara Bio). The library was sequenced single-end, 1×75 bp on a NextSeq2000 (Illumina) and was sequenced to a targeted coverage of 25 million reads. RNA-seq data was uploaded to sRNAbench (sRNAtoolbox) for analysis (*28*).

A supernatant after the second centrifugation step was sent to Novogene for RNA extraction and smRNA library prep with rRNA depletion and 20 million reads.

### Antibody development

Six-week-old BALB/cJ mice (The Jackson Laboratory) were immunized intraperitoneally with 100 µg synthetic PAP amyloid fibrils emulsified with incomplete Freuds adjuvant and boosted 3 times with 50 µg. The mice were allowed to rest for a month. A boost was then given in PBS and four days later the mouse was sacrificed, and the spleen harvested for fusion of spleen cells with NS-1 myeloma cells using PEG 1500 (Roche) (*52*). Complete selective IMDM medium containing hypoxanthine-aminopterin-thymidine (HAT), 20% FCS, pen-strep, and glutamine and was gently added to the fused cell pellet and the cells were kept at 37 °C for 3-5 hours to recover before being resuspended and plated into 96-well plates, containing a macrophage feeder layer to support hybridoma growth. After the cells were allowed to grow for 10 days, the hybridoma supernatants were screened by ELISA on synthetic PAP amyloid fibrils.

### Dot blot

Synthetic monomer or amyloid fibrils were spotted onto a nitrocellulose membrane (Bio-rad) to evaluate immunoreactivity for PAP amyloid fibril antibodies. Dot plots were blocked with Intercept® Protein-Free Block Buffer (LI-COR) and incubated at 4 °C for 1 hour with primary antibody. Blot was washed in TBS-T, incubated with IRDye® 680RD donkey anti-mouse secondary antibody (LI-COR) for 1 hour at RT, and washed again. Gel was imaged on the Odyssey CLx (LI-COR).

### Cryo-EM sample freezing

3 µL of samples were pipetted on glow-discharged R1.2/1.3 300-mesh, Au, Carbon Quantifoil EM grids (Quantifoil Micro Tools), which were plunge frozen in liquid ethane using a Vitrobot Mark IV semi-automated plunge freezer (Thermo Scientific) with blot times ranging from 3 seconds to 7 seconds. Grids were clipped into AutoGrids (Thermo Scientific).

### Cryo-EM data collection

High-resolution images were acquired at Columbia University Zuckerman Institute’s Titan Krios microscope (Thermo Fisher) equipped with an advanced BioQuantum-K3 imaging system. All images were captured at 300 kV using a Gatan K3 Summit detector. The movies were recorded with a pixel size of 1.074 Å, employing a dose rate of 1.5 electrons *per* Å^2^ for each frame. The defocus range was set between -1.2 and -2.5 µm.

### Single particle analysis

Movie frames were gain-corrected, aligned, dose-weighted, and summed using MotionCor2 (*53*) implemented in RELION (*54*). Parameters for the contrast transfer function were estimated from each motion-corrected micrograph using Gctf (*55*). Particles were manually picked. Particles were extracted in RELION with a box size of 810 Å for 2D classification.

### Mass spectrometry

The sarkosyl-insoluble fraction of SF was incubated with 4M guanidinium hydrochloride to disaggregate the amyloid fibrils. In solution, digestion was performed using LysC and trypsin, followed by desalting and LC-MS/MS for protein identification and quantification. The data were processed by MaxQuant. Mass spectrometry data was searched against the Uniprot human protein database. Mass spectrometry was also performed on murine sperm and embryos. Samples were lyased with lyase buffer (50 mM Tris pH 7.5, 150 mM NaCl, 1% Triton-X100), centrifuged at max speed, and supernatant was used for LC-MS/MS. The data was searched against the Uniprot human and mouse protein databases.

### Cryo-ET sample preparation, freezing, and cryo-CLEM

Bovine sperm (ABS Global) 250 cc straws were thawed in 37 °C water bath for 30 seconds and centrifuged for 10 minutes at 400 × *g*. Supernatant was removed, and sperm were re-suspended and washed with Human Tubule Fluid (HTF). HTF solution was made with 97.8 mM NaCl, 5 mM KCl, 0.2 mM MgSO_4_•2H_2_O, 28 mM HEPES, 20 mM sodium lactate solution (60% w/w), 0.4 mM sodium pyruvate, and 3 mM glucose at pH 7.4. At the last wash, supernatant was removed, and sperm were re-suspended in HBSS and 50 µg/mL PAP amyloid fibrils linked to gold nanobeads (nanopartz). The gold-nanobeads were linked to the fibrils according to the manufacturer’s protocol. Sperm incubated with the fibrils for 1 hour at 37 °C, washed, and re-suspended in HEPES buffer (10 mM HEPES, 150 mM NaCl, 10% BSA, pH 7.4). SYBR14 and propidium iodide were added to sperm according to the LIVE/DEAD Sperm Viability Kit protocol (Thermo Fisher Scientific). Live cells were sorted using FACS.

For cryo-ET sample preparation, 3.5 µl of live sperm were pipetted on glow-discharged R2/2, 200 hexagonal-mesh, Au, carbon Quantifoil EM grids (Quantifoil Micro Tools) which were plunge frozen in liquid ethane in a Vitrobot Mark IV semi-automated plunge freezer (Thermo Scientific) with a blot time 0.5 s, a wait time 15 s, and a blot force of -10. Grids were clipped into AutoGrids (Thermo Scientific) and placed in cryo-CLEM to determine ice thickness and quality, and to identify targets.

### Cryo-Focused Ion Beam-Scanning Electron Microscopy (cryoFIB-SEM)

To create lamellae, grids were loaded into the Fitzpatrick Lab Aquilos cryoFIB-SEM (Thermo Scientific) and eucentric height was taken on targets identified in cryo-CLEM. Grids were sputter coated with platinum metal and then coated with trimethyl(methylcyclopentadienyl)platinum(IV) with the onboard gas injection system (GIS). Trenches around the targets were milled at 40° to release tension and targets were then milled at angles between 7°-9° at 30 kV, using currents of 0.5 nA, 0.3 nA, 0.1 nA, and 50 pA for the focused ion beam. Lamellae were polished with a current of 10 pA for a final thickness between 150-200 nm.

### Cryo-ET data collection

A Titan Krios (Thermo Fisher) with a K3 camera, Gatan Imaging Filter system (Gatan), was used for data collection operating at 300 kV. All data was collected in super-resolution mode at a magnification of 53,000× with a pixel size of 1.7 Å/pixel. SerialEM (*56*) was used to set up and collect tilt series with total does of 120 e-/Å^2^, over 41 tilts comprised of 10 frames *per* tilt. Each tilt series was collected over a range of 120° in 3° increments following a dose-symmetric scheme, starting at a pre-tilt angle defined by the milling angle (7°-8°). A total of 74 lamellae were imaged.

### Tomogram reconstruction and segmentation

Tilt series were pre-processed in Warp 1.0.9 (*57*) where 2D CTF correction, Motion Correction, and creation of tilt stacks was performed. AreTomo (*58*) was used for tomogram alignment and reconstruction and denoised with Topaz 3D denoise (*59*). Membranes in tomograms were segmented with MemBrain (60) and Amira (Thermo Fisher), while amyloid fibrils linked to gold nanobeads were denoised in IsoNet (61) and segmented in Amira.

### High pressure freezing-freeze substitution serial section electron tomography

Flow sorted bovine sperm were fixed with freshly made fixative containing 2% paraformaldehyde, 2% glutaraldehyde in 0.1 M sodium cacodylate buffer (pH 7.2) on ice for 2 hours, washed with PBS, then loaded on 100 µm deep planchette hats filled with 20% bovine serum albumin (BSA)/PBS. The hats were coated with hexadecane, sealed in the planchette holder and high pressure frozen using Leica ICE high pressure freezer (Leica Microsystems). Freeze substitution was done using Leica EM AFS2 unit, all chemicals are purchased from Electron Microscopy Sciences, PA. Briefly, frozen hats were transferred into a mixture of 2% osmium tetroxide/0.1% uranyl acetate/98% acetone/2% ddH_2_O inside liquid nitrogen, and hold at -90 °C for 79 hrs. The samples were slowly warmed up to -60°C (5 °C/hr), hold 12 hrs. at -60 °C, then warm up to -30 °C (5 °C/hr) and hold for 6 hrs., then warmed up to 0 °C (6 °C/hr) for hold additional 6 hrs.

For morphology, the samples were rinsed in 100% acetone 2 × 1 hr at room temperature, removed from the hat, placed in EMbed812:acetone (1:1 ratio) for 1 hr., then EMbed 812:acetone (2:1 ratio) overnight. The samples were continually infiltrated in pure EMbed 2 × 2 hrs., flat embedded using two pieces of Aclar embedding film (Ted Pella, Inc. CA) which were situated and stabilized between two glass slides and polymerized at 60 °C for 48 hrs.

To enlarge the size of 3 nm gold nanobeads, we applied silver enhancement. In brief, frozen hats were transferred into a mixture of 2% glutaraldehyde/0.01% tannic acid/98% acetone at liquid nitrogen for freeze substitution as described above. After samples were warmed up to 0 °C, samples were washed with 100% ethanol, rehydration with 95%, 85%, 70%, 50% and 30% ethanol 10 minutes each step. The samples were washed with ddH_2_O, then underwent silver enhancement (HQ Silver enhancement kit, Nanoprobes, Yanphank, NY) in the dark for 10 minutes. The samples were continually washed with ddH_2_O, then underwent gold toning (2% NaAcetate, 3 × 5 min at RT; 0.05% HAuCl_4_, 10 min on ice; then 0.3% Na_2_S_2_O_3_, 2 × 10 min on ice) to prevent the oxidation of the silver. The samples were fixed in 2% OsO_4_ and 1.5% potassium ferrocyanide in phosphate buffer for 1.5 hours and *en bloc* stained with 1% uranyl acetate in aqueous solution overnight at 4 °C, then washed and dehydrated in a series of ice cold ethanol (30%, 50%, 70%, 85%, 95%, 100%, 100%), exchanged with room temperature acetone and embedded in EMbed812 resin (Electron Microscopy Sciences, EMS, PA) using Aclar embedding film.

200 nm serial sections were cut (Leica UC6 microtome) and collected on formvar coated slot cooper grids, stained with uranyl acetate and lead citrate by standard methods. Using a JEOL1400 Flash TEM, electron tomography tilt series were collected with a dual-axis tomography holder (Fischione 2040 dual-axis rotating holder) using SerialEM program for automated data collection (56) from -65° and +65° at 1.5-degree intervals. A second tilt series of the same area was collected after rotating the specimen support by ninety degrees.

### Isolation of human spermatozoa

Human sperm was isolated from cryopreserved semen (Lee Biosolutions) from anonymous, healthy donors. Frozen semen was thawed in water bath at 37 °C and then centrifuged at 400 × *g* for 5 minutes to pellet sperm. Supernatant was removed.

### FACS

Sperm cells were sorted after the use of the LIVE/DEAD Sperm Viability Kit protocol (Thermo Fisher Scientific). 300 µL of cells were strained using 5 mL Polystyrene Round-Bottom Tubes with Cell-Strainer Cap (Corning) and next analyzed and sorted using a Beckman Coulter MoFlo Astrios EQ (Beckman Coulter). Gated single cells were sub-gated and sorted on PI+/SYBR14+ cells for cryo-ET. For human spermatozoa, PI-(532-622)/SYBR14+(488-513) was sub-gated on AlexaFluor-350 (355-448) for analysis.

### IVF

Sperm preparation: C57BL/6J (The Jackson Laboratory) males were sacrificed, and sperm was harvested from the cauda epididymis and vas deferens. Sperm cells were washed with human tubule fluid (HTF) and incubated with synthetic PAP amyloid fibrils bound with ^13^C/^15^N labelled 2U-TAR RNA for 1 hour at 37 °C. The amyloid-RNA complex was washed and prepared as described in the sedimentation binding assay. After incubation, sperm cells were washed and resuspended in HTF. Sperm were also prepared similarly with EGFP mRNA (Trilink Bio Technologies).

Oocyte preparation: Female C57BL/6J mice, aged between 4 and 5 weeks, were super-ovulated by intraperitoneal (i.p.) injections of 5 international units (IU) mare serum gonadotropin. 48 hours later, 5 IU of human chorionic gonadotropin (hCG) were injected into females by i.p. 14-15 hours after hCG injection, females were sacrificed, and oocytes were collected.

Sperm and oocytes incubated together in HTF/GSH media in a petri dish for 30 minutes at 37 °C. If cumulus cells did not fall off, more sperm was added. After 3-4 hours of incubation, embryos were washed in HTF and incubated overnight. The next day, embryos at the 2-cell stage were collected and RNA (<200 nt) was extracted using a RNeasy Micro Kit (Qiagen) and enriched with biotin-TAT(47–57) (Anaspec) and streptavidin magnetic beads (NEB).

RNA sample preparation for mass spectrometry was performed as previously described by the Sidoli lab (62, 63). For each experimental sample, 2 µg of RNA were subjected to hydrolysis using the Nucleoside Digestion Mix (New England BioLabs, Cat # M0649S), following manufacturer instructions. Nano-liquid chromatography was configured with a two-column system consisting of a 300 μm ID × 0.5 cm C18 trap column (Dionex) and a 75 μm ID × 25 cm Reprosil-Pur C18-AQ (3 μm; Dr. Maisch GmbH, Germany) analytical nano-column packed in-house using a Dionex RSLC Ultimate 3000 (Thermo Scientific, San Jose, CA, USA). Chromatography was coupled online to an Orbitrap Fusion Lumos mass spectrometer (Thermo Scientific). The spray voltage was set to 2.3 kV and the temperature of the heated capillary was set to 275 °C. The full scan range was 110–600 m/z acquired in the Orbitrap at a resolution of 120,000. The source fragmentation energy was set at 30 V and RF lens % was set at 50. Signals of the nucleosides were extracted manually using the Xcalibur software.

### Statistical anlysis

Welch’s *t*-test was used to assess significance for the kinetics half-time plots (Fig. 2B and fig. S6D). Tukey’s multiple comparisons test was applied to evaluate differences between the means of % internalized sperm (Fig. 5A). An unpaired *t*-test was used to compare the means of heavy/light ratios from mass spectrometry analysis of TAR RNA (Fig. 5E).

---

**Fig. S1.**
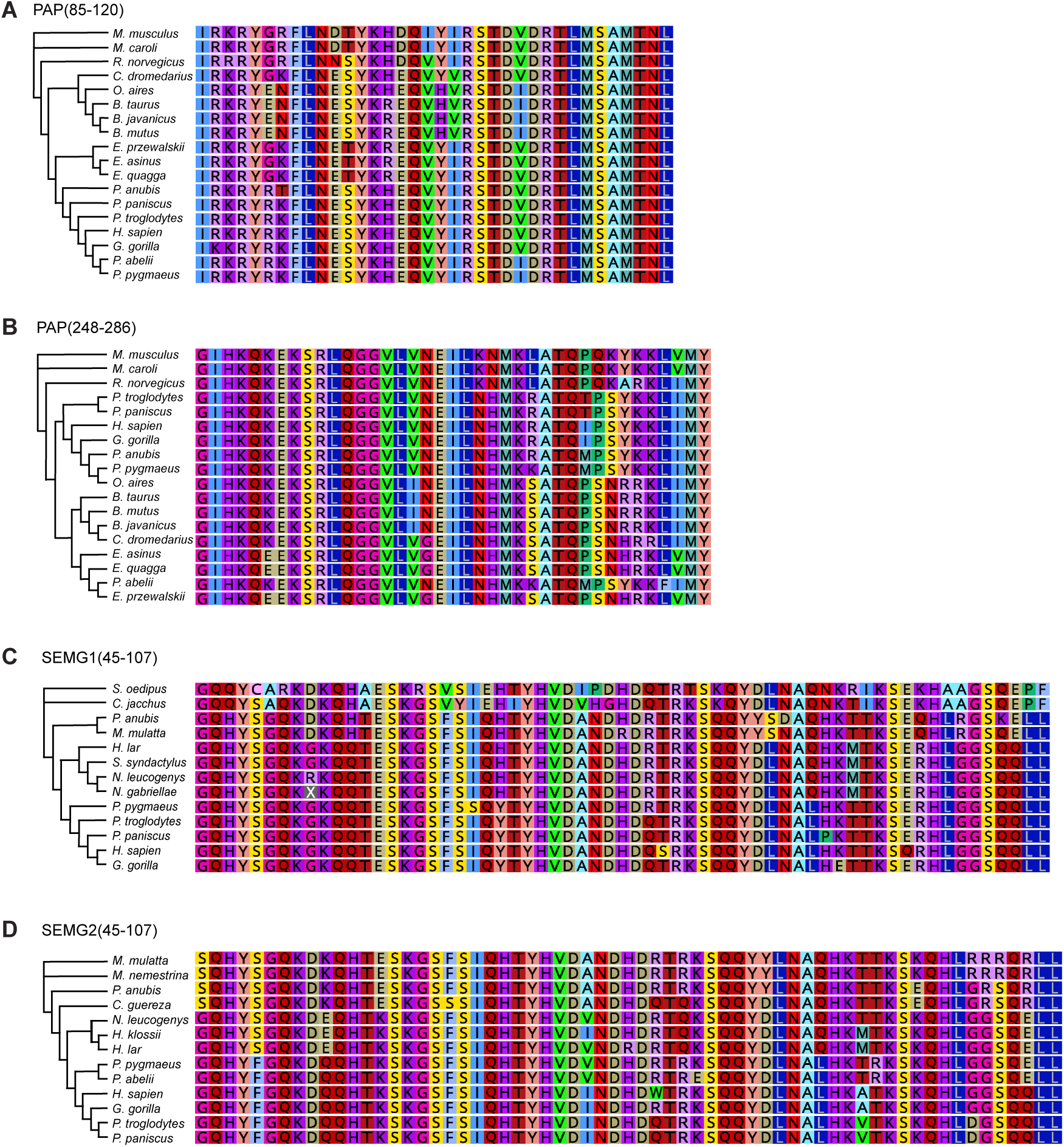
Phylogenetic relationships of seminal fluid peptide homologs across mammals. **(A-D)** Amino acid sequences of seminal fluid peptides from representative mammalian species were aligned and used to generate phylogenetic trees. Sequence homologs were identified using NCBI BLAST, and multiple sequence alignments and phylogenetic analysis were performed using Geneious.

**Fig. S2.**
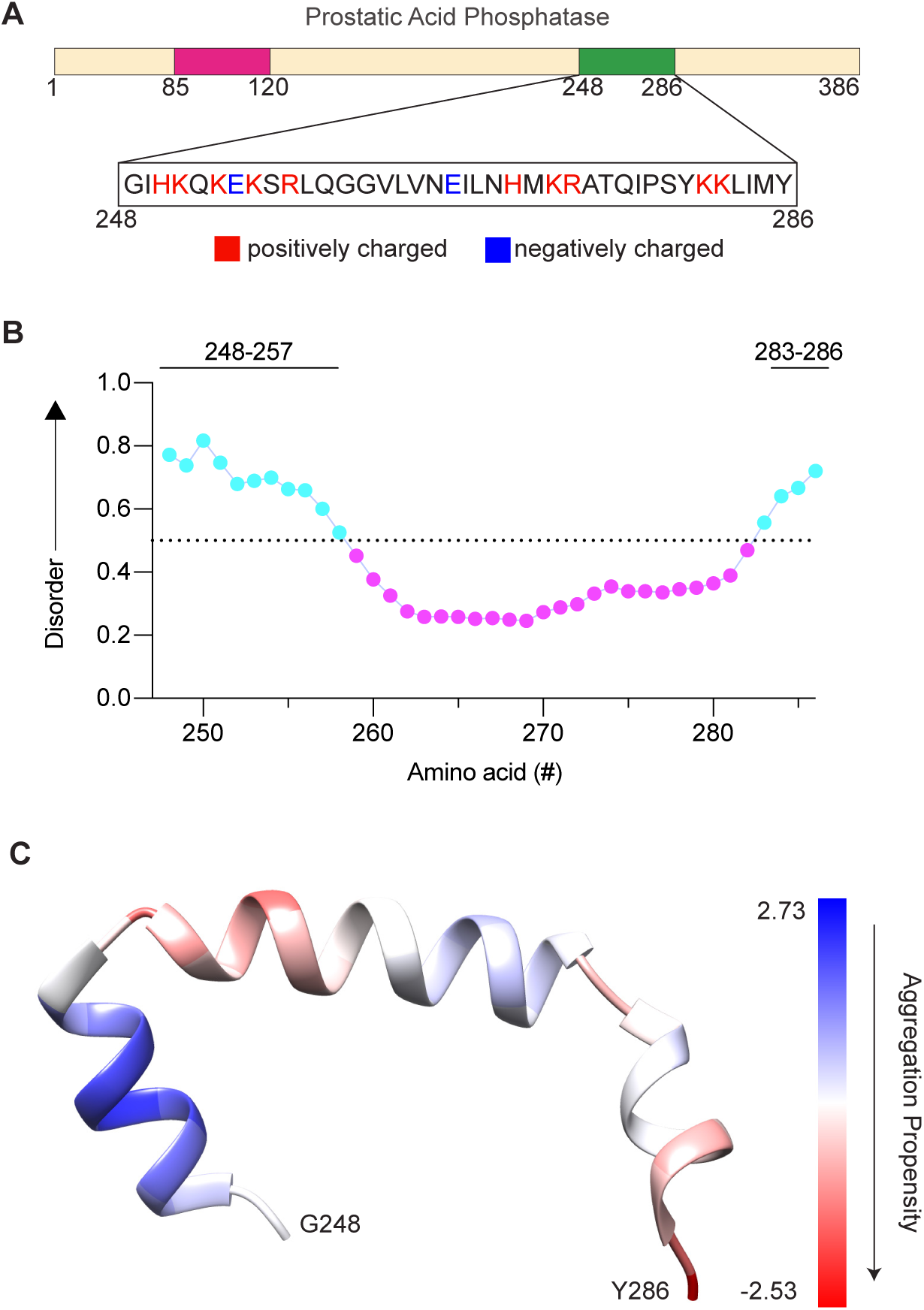
Sequence context and biophysical features of the PAP(248-286) region. **(A)** Location of the PAP(248-286) fragment within full-length prostatic acid phosphatase. **(B)** Intrinsic disorder propensity of PAP predicted using VSL2 (PONDR) (*19*); black solid lines denote predicted intrinsically disordered regions. **(C)** Predicted monomeric structure generated by AlphaFold3 (*64*), with aggregation propensity scored using CamSol (*20*).

**Fig. S3.**
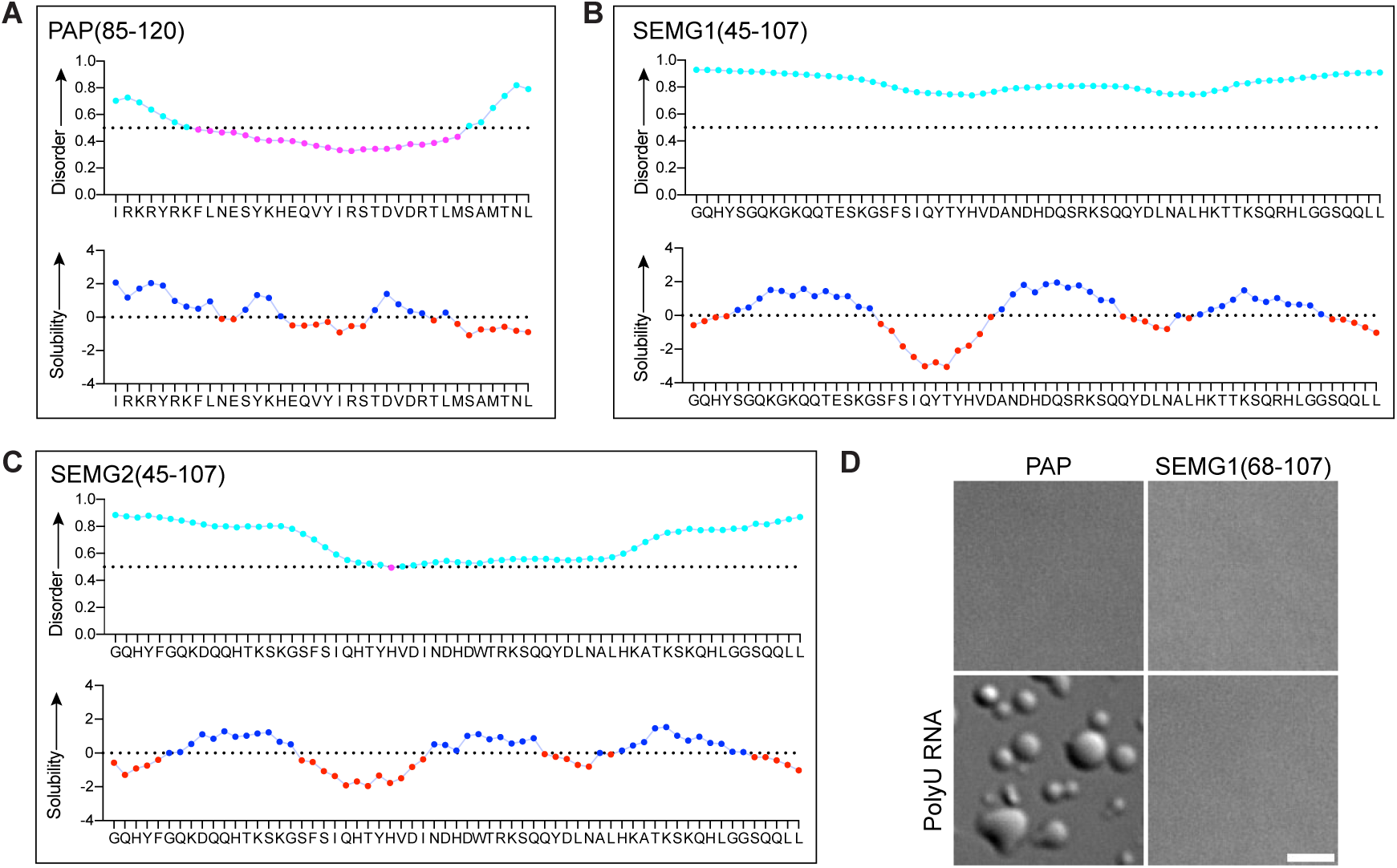
Intrinsically disordered and aggregation-prone regions in seminal fluid peptides. **(A-C)** Amino acid sequences of seminal fluid peptides with predicted intrinsic disorder (VSL2/PONDR) and solubility/aggregation propensity (CamSol). **(D)** Confocal images showing liquid-liquid phase separation (LLPS) of PAP or SEMG1(68-107) in the presence or absence of polyU RNA (500 µg/mL). Scale bar, 5 µm.

**Fig. S4.**
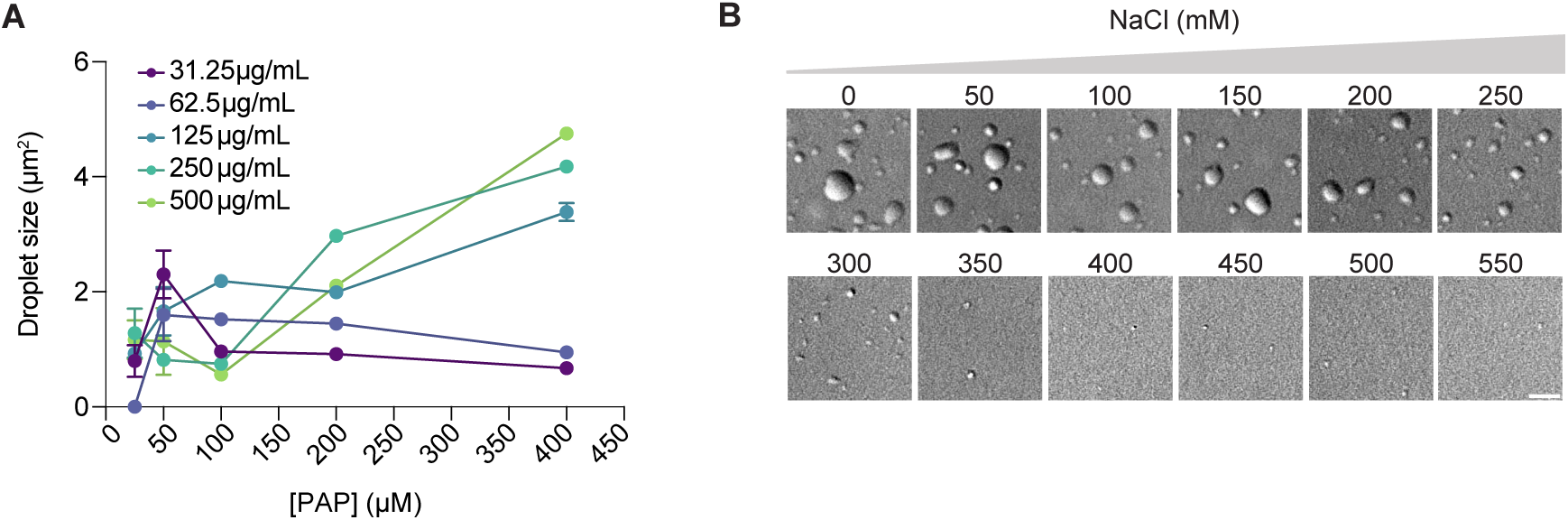
PAP undergoes LLPS with RNA in varying ionic conditions. **(A)** Quantification of PAP-polyU RNA droplet cross-sectional areas (µm^2^) from three independent experiments. **(B)** Confocal images of PAP (400 µM) mixed with polyU RNA (500 µg/mL) across a NaCl concentration gradient. Scale bar, 5 µm. Representative of *n* = 3 independent experiments.

**Fig. S5.**
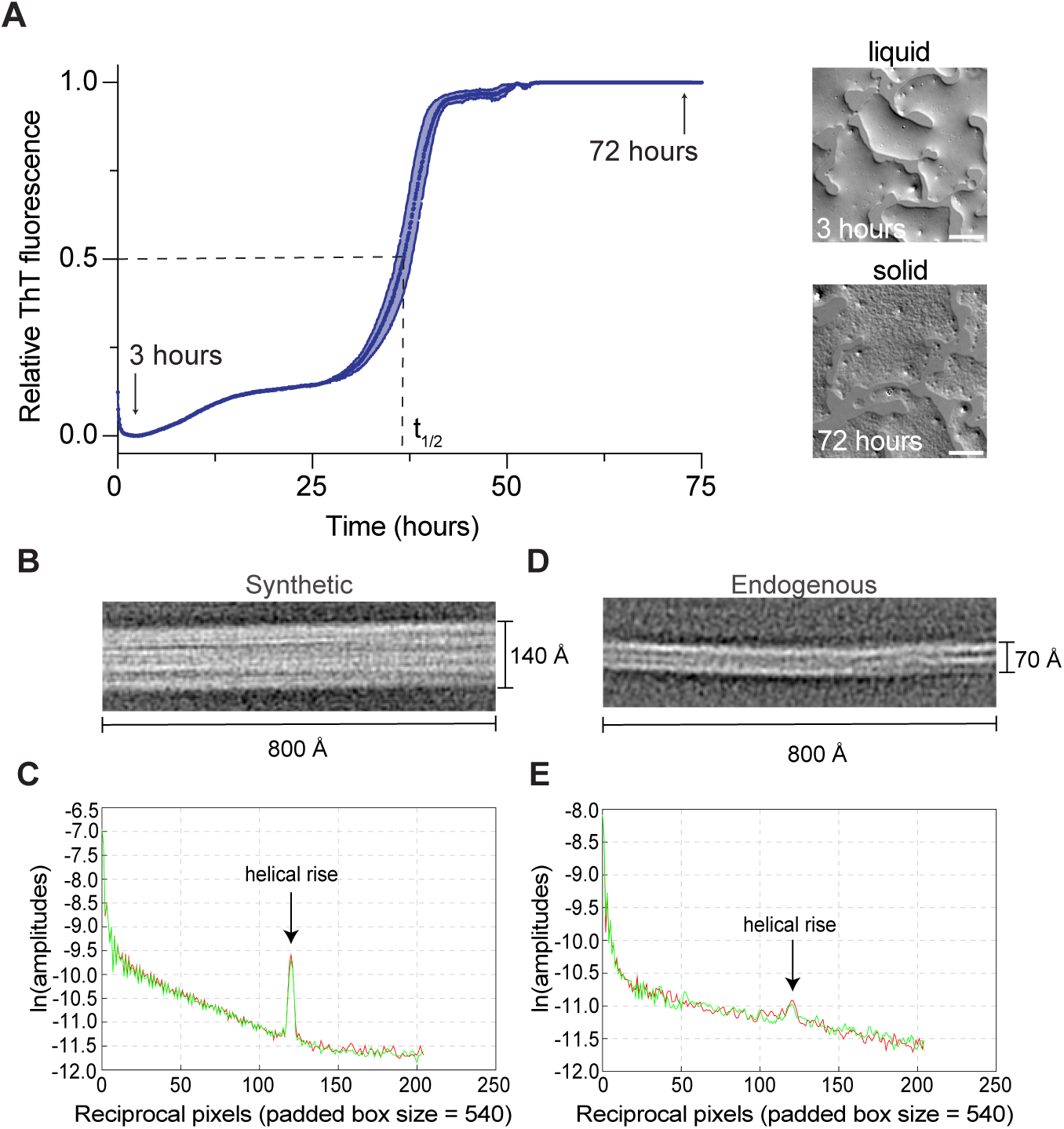
RNA accelerates PAP amyloid fibril formation. **(A)** ThT fluorescence kinetics of PAP (2 mg/mL) ± 30-nt polyU RNA-ATTO488 (60 µg/mL) at room temperature. Shaded region represents the SEM (*n* = 3). Confocal images from a parallel plate at 3 hours and 72 hours. Scale bars, 5 µm. **(B)** Representative 2D class averages of synthetic PAP amyloid fibrils. **(C)** Helical layer line analysis of synthetic PAP amyloid fibrils showing a helical rise of 4.7 Å consistent with β-strand spacing. **(D)** Representative 2D class averages of endogenous amyloid fibrils isolated from human seminal fluid. **(E)** Helical layer line analysis of endogenous amyloid fibrils showing the same 4.7 Å β-strand separation.

**Fig. S6.**
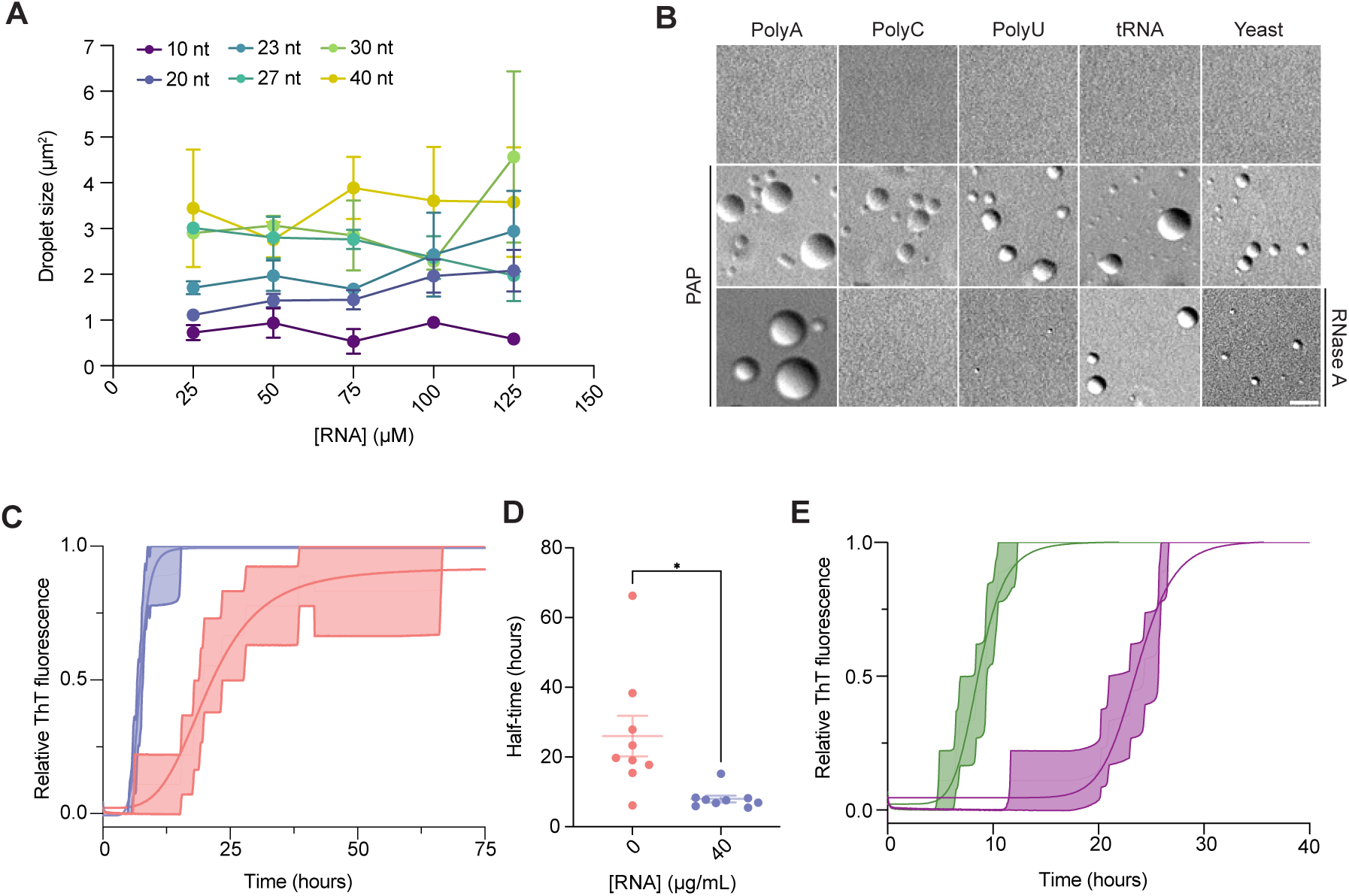
RNA length and concentration modulate PAP phase separation and fibrillization kinetics. **(A)** Quantification of PAP-polyU RNA droplet cross-sectional areas (µm²) from three independent experiments. **(B)** Confocal images of 500 µg/mL RNA alone, RNA + PAP (400 µM), or RNA + PAP + RNase A (500 µg/mL). Scale bar, 5 µm. Representative of *n* = 3 independent experiments. **(C)** ThT fluorescence kinetics of PAP (1 mg/mL) with (blue) or without (red) 40 µg/mL polyU RNA at 37 °C and 700 rpm. Lines show sigmoidal fits; shaded regions indicate SEM. Data reflects *n* = 3 biological × 3 technical replicates. **(D)** Half-time (t_1/2_) values derived from the curves in (C), indicating the time at which 50% of PAP has formed amyloid fibrils. *P* = 0.0149 (*). **(E)** ThT kinetics of PAP (1 mg/mL) with 23-nt (purple) or 40-nt (green) polyU RNA (30 µM). Lines and SEM shading as in (C). *n* = 3 biological × 3 technical replicates.

**Fig. S7.**
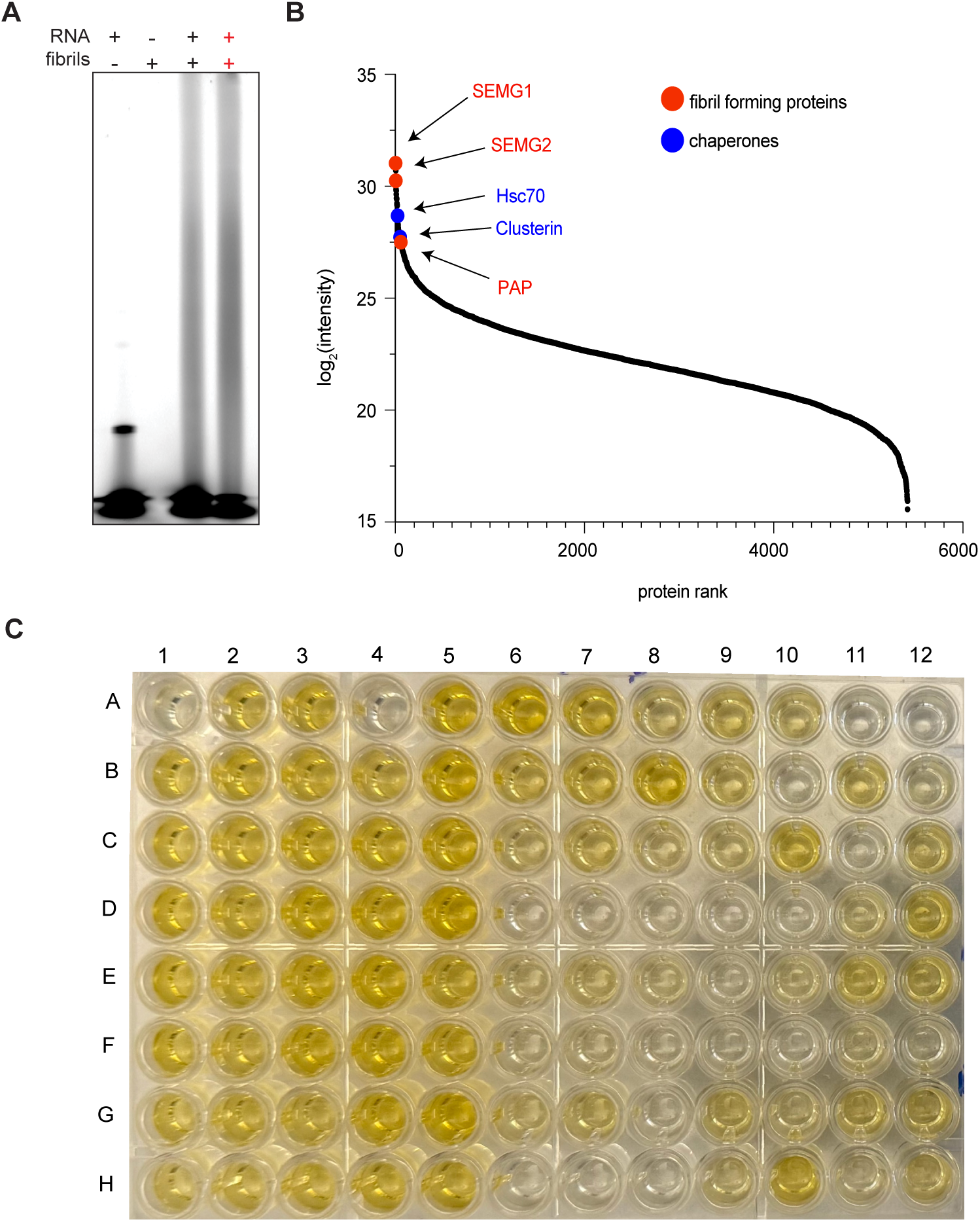
PAP amyloid fibrils bind RNA and are present with additional seminal fluid proteins. **(A)** Electrophoretic mobility shift assay (EMSA) showing that polyU RNA interacts with PAP amyloid fibrils. RNA remains associated with fibrils formed in the presence of RNA (red ++). **(B)** Proteins identified by mass spectrometry from the sarkosyl-insoluble fraction of human seminal fluid. PAP, SEMG1, and SEMG2 are among the most abundant components, and chaperones including Hsc70 and clusterin are also identified. **(C)** Selected hybridoma was single cell cloned twice. ELISA confirms clones of conformation-specific IgM antibody against synthetic PAP amyloid fibrils are positive.

**Fig. S8.**
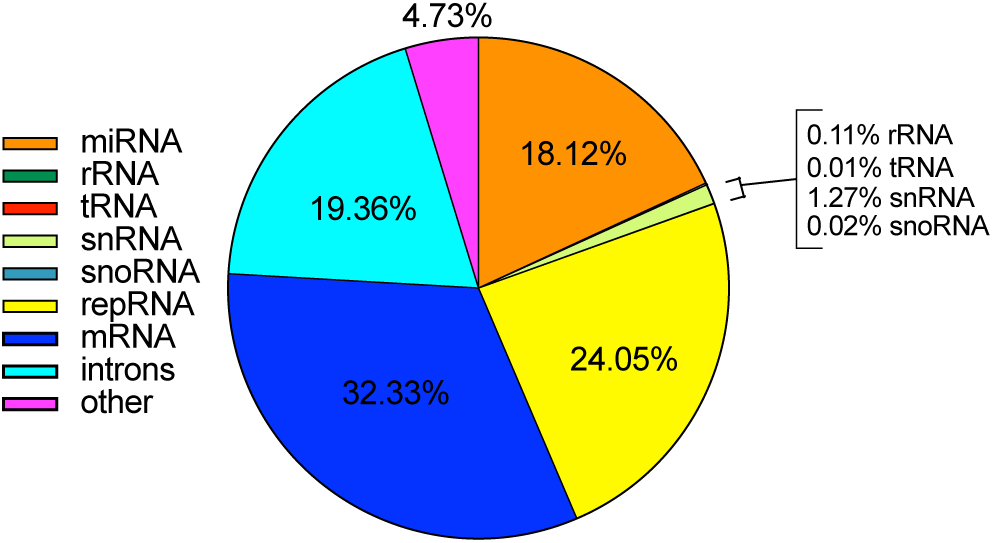
RNA-seq of endogenous seminal fluid. Percentage of mapped reads of the different RNA species detected in the supernatant of endogenous human seminal fluid.

**Fig. S9.**
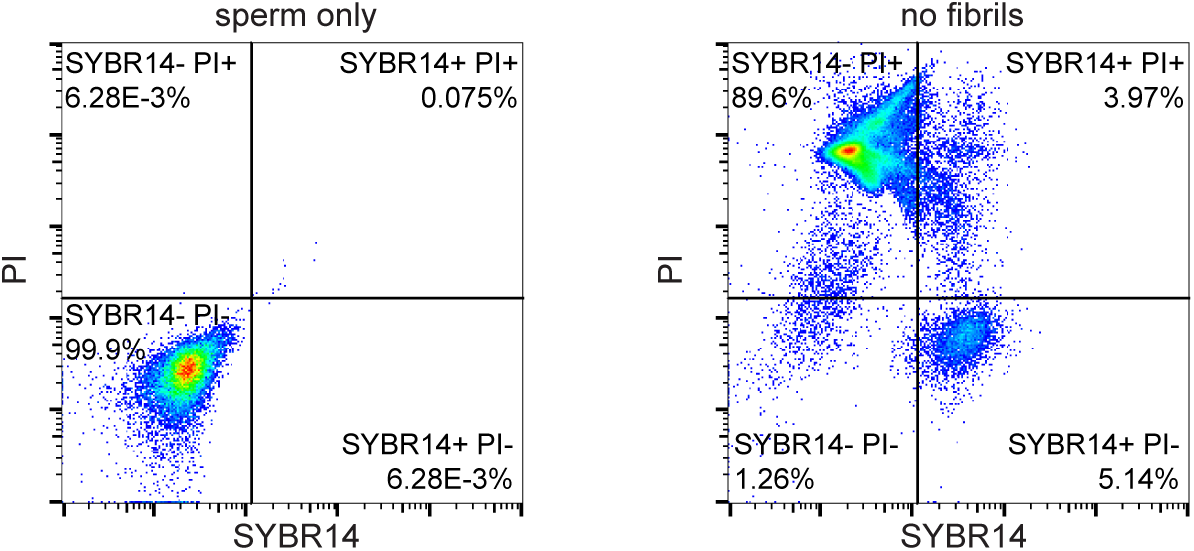
Flow cytometry control gating. Unstained (left) and fluorescently stained (right) bovine sperm were used to define gating boundaries in the absence of PAP amyloid fibrils.

**Fig. S10.**
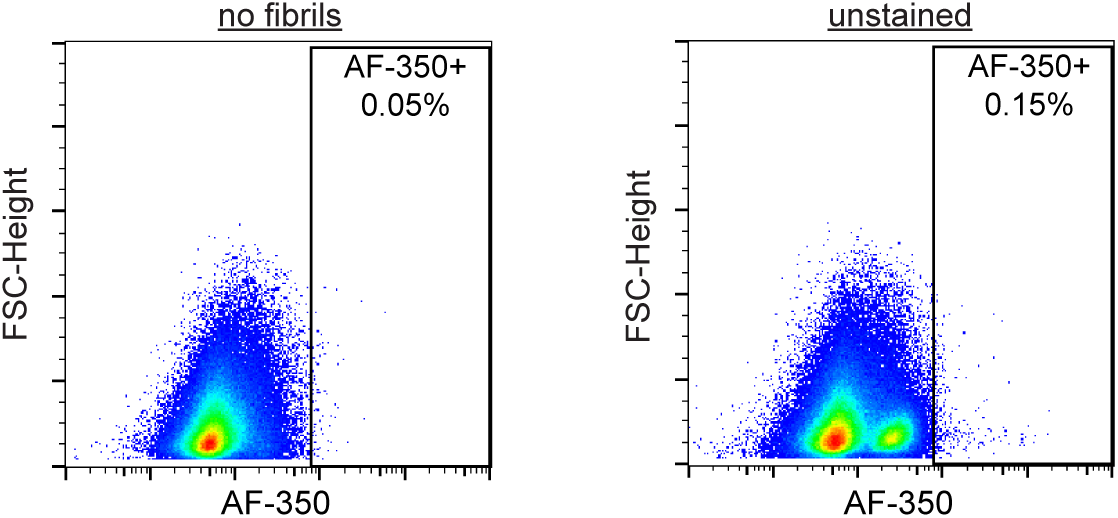
Trypsin reduces internalization of PAP amyloid fibrils by sperm. Flow cytometry (355-448) of human sperm incubated with no amyloid fibrils, unstained PAP amyloid fibrils, AF350-labeled PAP amyloid fibrils, or AF350-labeled PAP amyloid fibrils treated with trypsin.

**Fig. S11.**
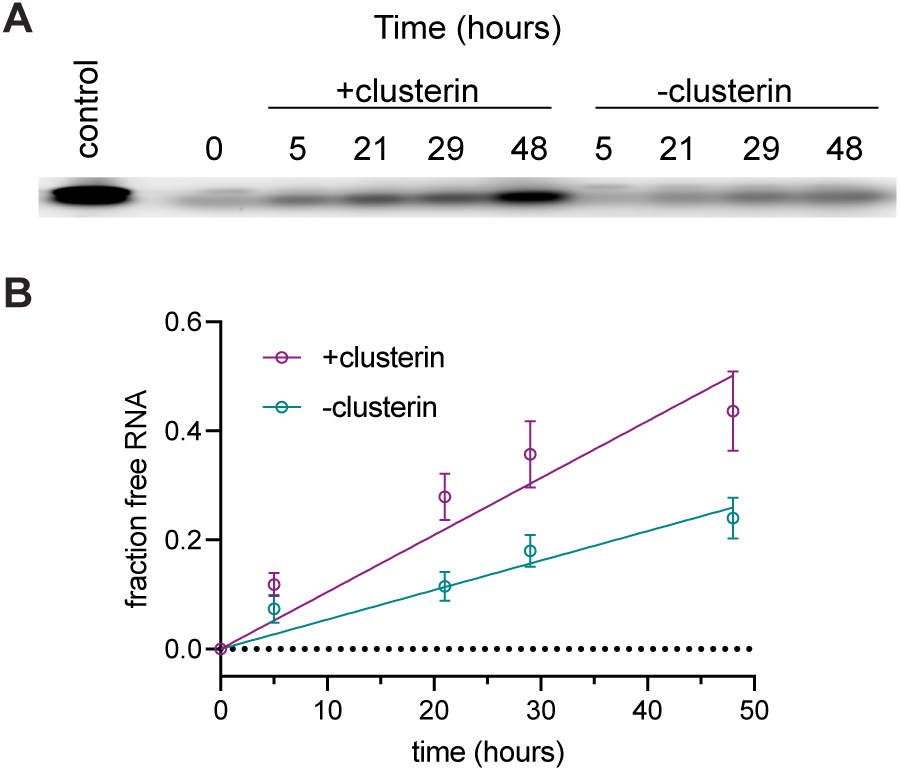
Clusterin increases the rate of RNA dissociation from PAP amyloid fibrils. **(A)** EMSA showing that RNA remains free in the presence of clusterin. *n* = 10 replicates. **(B)** Fraction of free RNA over 48 hours in the RNA dissociation assay. The dissociation rate (slope) in the presence of clusterin is 1.05 × 10^-2^ ± 8.17 × 10^-4^ fraction bound RNA *per* hour, compared to 5.40 × 10^-3^ ± 4.50 × 10^-4^ bound RNA *per* hour in the absence of clusterin.

**Fig. S12.**
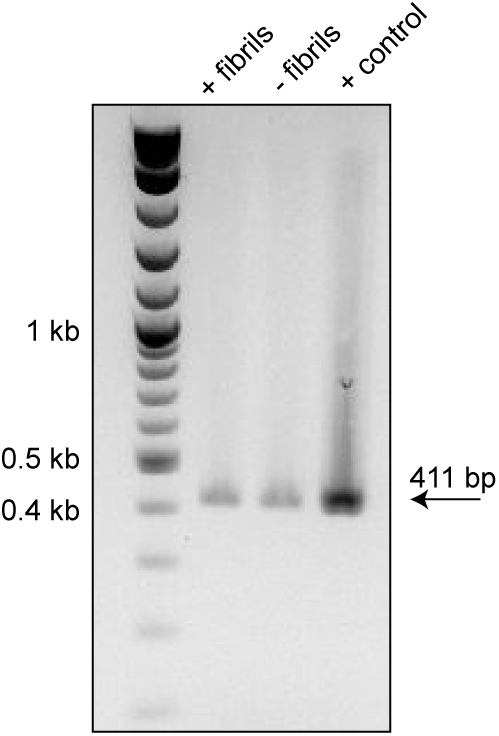
Detection of EGFP mRNA in 2-cell stage mouse embryos by RT-PCR. An amplicon of approximately 411 bp corresponding to the EGFP cDNA region amplified by nested primers is observed on an agarose gel. The positive control is purified EGFP mRNA.

**Table S1.** Human semen donor information for cryo-EM and mass spectrometry. Donor age, race, and health information is provided. The donors were pooled together and used for cryo-EM and mass spectrometry. They were all negative or non-reactive for anti-HIV 1 and 2, anti-HCV, and HBsAg by FDA approved methods.

| Donor | Birth Year | Gender | Race | Health Information |
| --- | --- | --- | --- | --- |
| 1 | 1999 | Male | Caucasian | None |
| 2 | 1967 | Male | Caucasian | None |
| 3 | 1966 | Male | Caucasian | None |
| 4 | 1986 | Male | Caucasian | None |
| 5 | 1995 | Male | Asian | None |
| 6 | 1990 | Male | Caucasian | None |
| 7 | 1970 | Male | Caucasian | None |
| 8 | 1966 | Male | Caucasian | High blood pressure |
| 9 | 1972 | Male | Caucasian | None |
| 10 | 1989 | Male | Caucasian | None |
| 11 | 1992 | Male | Caucasian | Depression/Anxiety, Allergies |
| 12 | 1981 | Male | Caucasian | Depression |
| 13 | 1981 | Male | Caucasian | None |

**Table S2.**
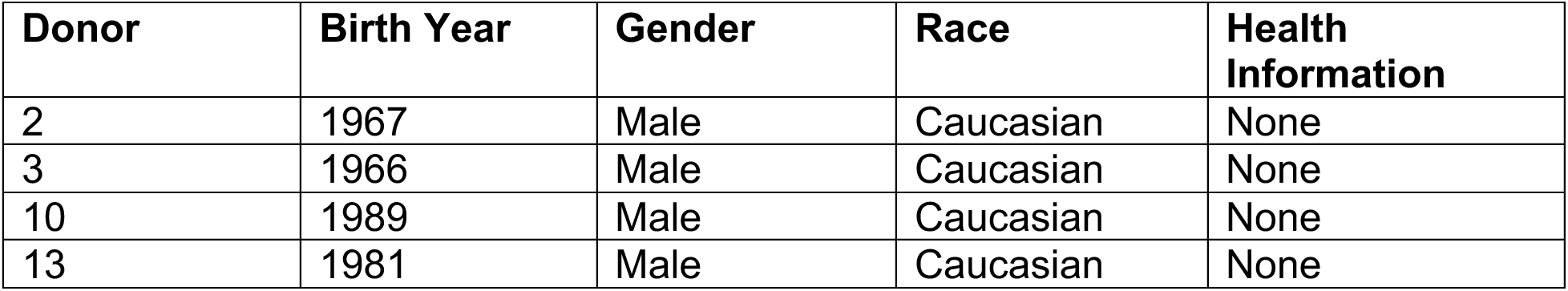
Human semen donor information for dot plot. Donor age, race, and health information is provided. The donors were pooled together and used for cryo-EM and mass spectrometry. They were all negative or non-reactive for anti-HIV 1 and 2, anti-HCV, and HBsAg by FDA approved methods.

| Donor | Birth Year | Gender | Race | Health Information |
| --- | --- | --- | --- | --- |
| 2 | 1967 | Male | Caucasian | None |
| 3 | 1966 | Male | Caucasian | None |
| 10 | 1989 | Male | Caucasian | None |
| 13 | 1981 | Male | Caucasian | None |

**Table S3.** Cryopreserved human sperm donor information for RNA-seq. Donor age, race, and health information is provided. The donors were pooled together and used for FACS. They were all negative or non-reactive for anti-HIV 1 and 2, anti-HCV, and HBsAg by FDA approved methods.

| **Donor** | **Birth Year** | **Gender** | **Race** | **Health Information** |
| --- | --- | --- | --- | --- |
| 1 | 1999 | Male | Caucasian | None |
| 3 | 1966 | Male | Caucasian | None |
| 14 | 1988 | Male | Asian/Caucasian | Hyperhidrosis |
| 15 | 1996 | Male | Caucasian | Anxiety/Depression |
| 19 | 2001 | Male | Caucasian | None |
| 20 | 1979 | Male | Caucasian | None |
| 21 | 1986 | Male | Caucasian | Colchicine,<br>Allopurinol |
| 22 | 1971 | Male | Caucasian | None |
| 23 | 1985 | Male | Caucasian | None |
| 24 | 1988 | Male | African<br>American | None |
| 25 | 1968 | Male | Caucasian | None |
| 26 | 1991 | Male | Caucasian | None |
| 27 | 1959 | Male | Caucasian | Verapamil |
| 28 | 1966 | Male | Asian | Metformin,<br>Atorvastatin |

**Table S4.** Cryopreserved human sperm donor information for FACS. Donor age, race, and health information is provided. The donors were pooled together and used for FACS. They were all negative or non-reactive for anti-HIV 1 and 2, anti-HCV, and HBsAg by FDA approved methods.

| **Donor** | **Birth Year** | **Gender** | **Race** | **Health Information** |
| --- | --- | --- | --- | --- |
| 8 | 1966 | Male | Caucasian | High blood pressure |
| 10 | 1989 | Male | Caucasian | None |
| 14 | 1988 | Male | Asian/Caucasian | Hyperhidrosis |
| 15 | 1996 | Male | Caucasian | Anxiety/Depression |
| 16 | 1992 | Male | Caucasian | None |
| 17 | 1996 | Male | Caucasian | None |
| 18 | 1983 | Male | Caucasian | None |

